# Visual experience shapes functional connectome gradients

**DOI:** 10.1101/2025.02.28.640746

**Authors:** Cemal Koba, Joan Falcó-Roget, Olivier Collignon, Katarzyna Rączy, Marina Bedny, Mengyu Tian, Marcin Szwed, Anna-Lena Stroh

**Author notes:** shared senior authorship (Marcin Szwed, Anna-Lena Stroh). **Author contributions Cemal Koba:** Conceptualization, Methodology, Software, Formal analysis, Data curation, Writing - Original Draft, Writing - Review & Editing, Visualization, **Joan Falco Roget:** Methodology, Writing - Review & Editing, **Olivier Collignon:** Funding acquisition, Data acquisition, Writing - Review & Editing, **Katarzyna Raczy:** Investigation, Data curation, Writing - Review & Editing, **Marina Bedny:** Funding acquisition, Data acquisition, Writing - Review & Editing, **Mengyu Tian:** data curation **Marcin Szwed:** Funding acquisition, Data acquisition, Writing - Review & Editing, Supervision **Anna-Lena Stroh:** Conceptualization, Methodology, Software, Formal analysis, Data curation, Writing - Original Draft, Writing - Review & Editing, Visualization, Supervision.

## Abstract

The human cortex is organized along continuous functional gradients that capture systematic transitions in functional connectivity across the brain. These gradients describe large-scale organizational principles, including hierarchical transitions from unimodal to transmodal regions. Here, we provide the first characterization of cortical gradients in a large sample of congenitally blind (n = 41) and sighted (n = 44) adults to assess the relative contributions of intrinsic (genetic) and experiential factors to cortical gradient organization. Using resting-state fMRI, we compared functional connectome gradients and their association with cortical structure. Both groups exhibited similar principal gradients: unimodal to transmodal, somatosensory to visual, and frontoparietal segregation, demonstrating that the fundamental scaffold of cortical organization emerges largely independently of visual experience. However, blindness altered specific features of the functional connectome: the visual network was more segregated from the sensorimotor network and more integrated with transmodal and frontoparietal networks. Moreover, blind individuals showed reduced canonical hierarchical ordering within early visual areas, weaker structure–function coupling in visual and temporal regions, and altered functional areal boundaries in V1. These findings suggest that the development of large-scale cortical gradients reflects a genetically guided scaffold that is subsequently refined by sensory experience.

## 1 Introduction

Processing in the brain is hierarchically organized, with sensory information flowing through successive stages of increasing complexity before reaching high-level cognitive areas^1^. Recent advances in data-driven analyses of resting-state functional magnetic resonance imaging (rs-fMRI) now make it feasible to study such functional hierarchies *in vivo*^2,3^. Gradient-based methods, in particular, reveal smooth spatial transitions in functional connectivity across the cortex, capturing key organizational principles of the brain, such as the progression from early sensory regions to transmodal association areas, or the differentiation between modality-specific systems^2^. The extent to which this hierarchical organization is shaped by sensory experience remains unclear. Studying people born blind can provide valuable insights into this issue, as it can reveal how the lack of sensory input into one sensory network changes the functional architecture not only within this network but across the broader cortex.

Prior studies in blind individuals have revealed both experience-dependent and -independent aspects of visual cortex organization. While the recruitment of early visual areas for higher-order processing^4,5^ and altered functional connectivity patterns^4–9^ may reflect changes in cortical hierarchy^10^, other findings suggest that the hierarchical organization is preserved. For example, blind individuals activate the fusiform face area when hearing voices^11^; the parahippocampal place area when processing environmental sounds^12^, and hMT+/V5 in response to auditory motion^13^. While these studies provide compelling evidence that the hierarchical organization is at least partially preserved in blind individuals, they generally offer isolated observations rather than a comprehensive view. By focusing on single regions or specific connections, they yield “point estimates” of reorganization rather than capturing the full extent of the cortical processing hierarchy. Thus, the question of how visual experience shapes the hierarchical organization of the visual system and its integration within large-scale brain networks remains unanswered.

Gradient-based analyses provide an opportunity to move beyond such “point estimates” by characterizing continuous patterns of connectivity across the whole cortex. Three principal gradients of functional connectivity have been identified^14^. The principal gradient (G1) extends from unimodal to transmodal areas and closely corresponds to classic models of the cortical hierarchy^15,16^, making it ideally suited to investigate the effects of sensory experience on the cortical hierarchy. The second gradient (G2) spans between the visual and sensorimotor systems^14^, differentiating regions by their involvement in modality-specific processing. The third gradient (G3) extends between areas of the brain that are typically involved in attentional modulation (fronto-parietal attention and control networks) and those that are involved in the sensory-motor and semantic content of representations (default mode, somatosensory/motor and visual networks)^17,18^. These gradients provide an ideal framework for studying how sensory experience influences the macroscale organization of the human brain.

The structural organization of the human cortex seems to follow similar macroscale gradients as the functional organization. A tight coupling between structure and function has been observed within primary sensory and motor areas, whereas in transmodal association areas, structure and function seem to diverge^19–22^. Strong coupling indicates that local structural features closely constrain functional architecture, whereas weaker coupling suggests greater functional flexibility or divergence from structural constraints. Although numerous studies have investigated both structural and functional plasticity in blind individuals, the relationship between the two remains largely underexplored, particularly in areas beyond the visual cortex^23,24^. Exploring these relationships may provide new insights into how cortical structure adapts to support the functions associated with the visual cortices of blind individuals.

Functional gradients are valuable not only for studying the relationships between different cortical networks but also for mapping the functional organization within networks^25,26^. Previous studies have used this approach to study the hierarchical organization within multiple brain regions^25,27–29^ and revealed gradients that show a strong correspondence with canonical functional networks and corresponding behavioral domains^28–31^, geometric distance^30^, cortical morphology^30^ and microstructure^27^, as well as gene expression^31^. Moreover, previous studies have used resting-state functional connectivity gradients for cortical area parcellations exhibiting strong correspondence with task activation maps and with probabilistic V1 and V2^32,33^. These functional gradients therefore serve as an effective data-driven tool to parcellate functional regions in the early visual cortex.

To summarise, we use data-driven gradient-based analyses to analyse rs-fMRI data acquired from a large sample of congenitally blind and sighted individuals, to test how visual experience shapes functional cortical gradients and their alignment with anatomy. First, we compared macroscale functional connectivity gradients between congenitally blind and sighted individuals to move beyond the “point estimates” of plasticity in previous studies. Then, we investigated how sensory experience shapes the functional organization *within* the visual system by comparing connectopic maps of the visual system between blind and sighted individuals. Finally, we test how visual experience shapes structure-function coupling by measuring alignment of functional connectivity gradients and structural features across the cortex in congenitally blind and sighted individuals. To this end, we developed a novel approach, where multiple macroanatomical measures (thickness, curvature, sulcal depth, surface area, and volume) are combined via principal component analysis to derive a single structural index for each participant and structure-function coupling is quantified by correlating this structural profile with large-scale functional gradients in blind and sighted individuals. Together, this work aims to elucidate how visual experience shapes the brain’s macroscale functional organization and its alignment with underlying structure.

## 2 Methods

### 2.1 Datasets

#### 2.1.1 Dataset 1

Complete details of the dataset and imaging parameters are given in Pelland et al.^34^ The entire dataset included 50 individuals who participated in a single five-minute functional MRI run (136 volumes). Participants were instructed to keep their eyes closed, relax, and not think about anything in particular. Functional time series were acquired using a 3-T TRIO TIM (Siemens) equipped with a 12-channel head coil. Multislice T2*-weighted fMRI images were obtained with a gradient echo-planar sequence using axial slice orientation; repetition time (TR) 2200 ms; echo time (TE) 30 ms; functional anisotropy (FA) 90 degrees; 35 transverse slices; 3.2 mm slice thickness; 0.8 mm gap; field of view (FoV) 192 × 192 mm^2^; matrix size 64 × 64 × 35; voxel size 3 × 3 × 3.2 mm^3^. A structural T1-weighted 3D magnetization prepared rapid gradient echo sequence (voxel size 1 × 1 × 1.2 mm^3^; matrix size 240 × 256; TR 2300 ms; TE 2.91 ms; TI 900 ms; FoV 256; 160 slices) was also acquired for all participants. All of the procedures were approved by the Research Ethic and Scientific Boards of the Centre for Interdisciplinary Research in Rehabilitation of Greater Montreal and the Quebec Bio-Imaging Network. Experiments were undertaken with the understanding and written consent of each participant.

The dataset consisted of 50 participants, including both congenitally blind and late blind participants and the respective sighted matches. In this study, we focused on the group of congenitally blind individuals and their sighted controls. The final sample thus included 28 participants, including 14 congenitally blind individuals (9 males, 5 females, mean age 43.93 ± 11.19 years, 12 right-handed, 2 ambidextrous), and 14 sighted controls, who were matched for age and sex with the sample of congenitally blind individuals (sighted controls: mean age 40.92±13.64 years, 7 males, 7 females, 13 right-handed, 1 left-handed). The dataset was made available in the Brain Imaging Data Structure (BIDS^35^) format. Along with the folder and naming standardization, the first four volumes of functional runs were removed to avoid stabilization artifacts (leaving 132 volumes in total), and the resolution of the functional runs was interpolated to 3 × 3 × 4 mm^3^. The data providers confirmed that no other preprocessing steps were applied during the standardization procedure.

#### 2.1.2 Dataset 2

Dataset 2 was shared by Bedny and Tian^36^, with a comprehensive description available in Tian et al.^37^. MRI anatomical and functional images were collected on a 3T Phillips scanner at the F. M. Kirby Research Center. T1-weighted anatomical images were collected using a magnetization-prepared rapid gradient-echo (MP-RAGE) in 150 axial slices with 1 mm isotropic voxels. Resting-state fMRI data were collected in 36 sequential ascending axial slices for 8 minutes. TR = 2 s, TE = 0.03 s, flip angle = 70°, voxel size = 2.4 × 2.4 × 2.5 mm, inter-slice gap = 0.5 mm, (FoV) = 192 × 172.8 × 107.5 mm^3^. Participants completed 1 to 4 scans of 240 volumes each (average scan time = 710.4 seconds per person). During the resting state scan, participants were instructed to relax but remain awake. Sighted participants wore light-excluding blindfolds to equalize the light conditions across the groups during the scans.

50 sighted adults and 30 congenitally blind adults contributed the resting state data (sighted: n = 50; 30 females; mean age = 35.33 ± 14.65; mean years of education = 17.08 ± 3.1; blind: n = 30; 19 females; mean age = 44.23 ± 16.41; mean years of education = 17.08 ± 2.11. Blind and sighted participants had no known cognitive or neurological disabilities (screened through self-report). A board-certified radiologist read all adult anatomical images and no gross neurological abnormalities were found. All the blind participants had at most minimal light perception from birth. Blindness was caused by pathology anterior to the optic chiasm (i.e., not due to brain damage). All participants gave written informed consent under a protocol approved by the Institutional Review Board of Johns Hopkins University. The dataset was made publicly available online after normalizing anatomical and structural data to MNI space, changing the voxel resolution of T1 images to 2 mm^3^ and functional images to 2.4 × 2.4 × 3 mm.

#### 2.1.3 Dataset 3

The dataset was collected in Malopolskie Centrum Biotechnologii, Jagiellonian University, Kraków on a 3T Siemens Magnetom scanner. T1-weighted anatomical images were collected using a magnetization-prepared rapid gradient-echo (MP-RAGE) in 1 × 1 × 1 mm voxels. Resting-state fMRI data were collected for 15 minutes. TR = 1.5 s, TE = 0.027 s, flip angle = 90°, voxel size = 3 × 3 × 3.5 mm. During the resting state scan, participants were instructed to relax but remain awake. The study was approved by the ethics committee of the Jagiellonian University. Written informed consent was obtained from all participants before the experiment. Participants were reimbursed for taking part in the study. The participants were screened and had no disability apart from vision loss. The dataset includes 17 congenitally blind participants (mean age: 25.75 ± 5.72, 6 females, 11 males), and is organized in BIDS format.

#### 2.1.4 Dataset curation and quality check

In total, 103 participants were retained across the three datasets following data organization and visual quality control. After an initial visual inspection, scans exhibiting artifacts—most commonly motion-related blurriness or incomplete slices—were excluded, resulting in 58 participants (36 sighted and 22 congenitally blind) out of the original 80. Datasets 1 and 3 required no exclusions, and their full samples of 28 (14 sighted and 14 congenitally blind) and 17 congenitally blind participants, respectively, were included in subsequent preprocessing steps.

### 2.2 Preprocessing

The datasets were preprocessed with Micapipe^38^, an automatic processing pipeline using state-of-the-art software for processing structural and functional MRI data such as AFNI^39^, FSL^40^, ANTs^41^, FreeSurfer^42^, FastSurfer^43^, and Connectome Workbench (RRID:SCR_008750). Full details of the analysis pipeline can be found at https://micapipe.readthedocs.io/. Briefly, each T1-weighted run was LPI-reoriented, deobliqued, and oriented to standard space (MNI152), a bias-field correction was applied, and the intensity of the images was rescaled between 0-100. Whole-brain, gray matter (GM), white matter (WM), and cerebrospinal fluid (CSF) segmentation was performed. The T1 image was then non-linearly registered to MNI152. Surface meshes in individual spaces were created. Affine registration from fsaverage5 mesh to individual meshes were calculated. Using these registration matrices, the Glasser parcellation^44^, which is available in the Micapipe library for the fsaverage5 mesh, was sent to subject space. Using the 360 available regions of interest (ROI) in the Glasser parcellation, regions-wise statistics such as cortical thickness, mean curvature, surface area, sulcal depth, and cortical volume were calculated.

Functional images were reoriented to LPI orientation, and a motion-correction algorithm with 6 parameters (rigid-body) was applied to the time series. The motion-corrected time series were then projected to the fsnative5 mesh and smoothed with a 10 mm Gaussian kernel. The outputs from Micapipe were further cleaned from nuisance regressors using a custom script written in MATLAB. The 36P strategy without global signal regression, described in Satterthwaite et al.^45^, was adopted to choose the regressors of no interest. Specifically, 6 motion regressors, mean signal from white matter and CSF, their first derivatives, power, and power of the first derivatives were chosen. In addition, mean framewise displacement and a linear trend were added to the model. These regressors were removed from the fMRI data using a multiple linear regression model and the residuals were saved. A band-pass filter between 0.01-0.1 was applied to the residuals.

### 2.3 Calculating the functional connectivity gradients

For each participant, preprocessed time series on fsaverage5 mesh were parcellated using the Glasser atlas (fsaverage5 version available in the Micapipe library). The averaged time series were then correlated with each other, resulting in 360 × 360 correlation matrices for each participant. The matrices were normalized via Fisher’s Z transformation.

Functional connectivity gradients based on normalized correlation matrices were generated as described in Margulies et al.^14^ (2016) using the available functions in the BrainSpace MATLAB toolbox^46^. In short, correlation matrices were thresholded to keep the top 10% of the connections. Cosine similarity of the sparse matrices was calculated. A non-linear dimension reduction technique, diffusion mapping, was applied to the similarity matrices with an alpha value of 0.5 for the manifold. This step generated 10 gradient values that explain the functional connectivity profile of each ROI and for each participant. Lastly, Procrustes alignment was used on the gradients to make them comparable. Since diffusion mapping can result in values in the opposite directions, a reference gradient available in BrainStat library^47^ was used for alignment, which was calculated with the same parameters on fsaverage5 mesh. This procedure generated “gradient scores” that show the position of each region in the gradient space for the first 10 gradients.

To decrease noise, an inter-subject correlation matrix was calculated based on scores of the first gradient. Participants who showed a smaller Pearson’s r than the absolute value of 0.4 were removed from further analyses. The final sample for the group comparisons included 44 sighted and 41 congenitally blind participants.

### 2.4 Statistical analyses

#### 2.4.1 Connectivity gradients across neocortex

After removing the effect of age and sex from the gradient scores via a multiple linear regression model, the residuals were used to compare the differences across the groups of sighted and blind individuals for each ROI via two-sample t-tests. This comparison was repeated for the first three gradients, independently. The results were corrected for multiple comparisons with the Benjamini-Hochberg false-discovery rate (FDR) procedure at p = .05. Following the region-wise comparison of the gradient scores, the gradient range (the maximum value minus the minimum value) was compared between groups using a multiple linear regression model where age and sex were included as regressors of no interest. In addition, the explained variances of the gradients and the variance of the gradient scores were compared between groups of blind and sighted individuals with a two-sample t-test. Effects of age and sex on the gradient range, variance, and explained variance were accounted for using a linear regression before applying a t-test.

#### 2.4.2. Community based profiles

To summarise the findings of the ROI analyses, we conducted a network-based z-score analysis of gradient scores for each of the first three gradients. To this end, mean gradient scores of each network were compared between the groups of blind and sighted individuals with two-sample t-tests with multiple comparison correction at a false discovery level of 0.05 (pFDR < 0.05). In addition, we assessed potential differences between the groups in the functional distance between established functional networks^48^. Between-network functional distance was calculated as the distance between network medians. Given that we expected that most of the changes between the two groups would be observed with respect to the visual network, we focused our analyses on the differences in functional distance between the visual network and the rest of the networks.

We used R^49^ and afex^50^ to perform a linear mixed effects analysis on the network distances. Separate models were run for each of the three first gradients. As fixed effects we entered group (blind vs sighted) and network (somatosensory/motor, dorsal attention, ventral attention, limbic, frontoparietal, and default mode network), as well as their interaction terms into the linear mixed model (LMM). In addition, age and sex were added as fixed effects into the model. As random effects, we had intercepts for participants. P-values for fixed effects were obtained with the Satterthwaite method and were considered significant at p < .05. Post-hoc comparisons of significant interactions were conducted using approximate z-tests on the estimated marginal means (EMMs) using the emmeans package^51^. The resulting p-values were corrected for multiple comparisons following the procedure proposed by Holm (1979).

#### 2.4.3 Representation of the early visual areas in the gradient space

To test whether the functional organization in the gradient space reflects the hierarchy of the visual areas, and whether this representation is different in the blind group, the position of the early visual areas within the first three gradients was checked in terms of their ranking.

Our first aim was to see if the median ranks of the early visual areas reflect the functional hierarchy from V1 to V4 in the sighted and blind groups. Group-level median scores were calculated using the mean gradient scores over the four early visual areas for both groups, and the median ranks were ordered from lowest to highest, and their ranking in this order (1 for the lowest value, 4 for the highest value). For each early visual area, a number between 1 and 4 was appointed after this step depending on their gradient score. In order to validate the group-level median ranks, a bootstrap resampling approach was employed. 10000 bootstrap replicates were generated by randomly sampling participants with replacement. For each bootstrap sample, the median values for each visual region for sighted and blind groups were computed. The ranks of median values were then sorted in ascending order, with ranks assigned from 1 (lowest median) to 4 (highest median). These rank combinations were recorded for each bootstrap replicate. The frequency of unique rank patterns (e.g., “V1-V2-V3-V4”) was counted to determine the most common configurations across bootstrap samples. This approach revealed the distribution of the rank order patterns across the bootstrap sample.

The bootstrap approach highlights the variance across the resampled sample’s medians. To be more specific to our sample, subject-specific rank orders were also calculated. The frequency distribution of 24 possible rankings was compared via a Chi-square test for independence. Observed counts were compared with the expected counts under the null hypothesis that both groups exhibited the same distribution of ranking orders. This analysis was useful to compare the overall ranking order frequency between groups. To test the difference of a specific ranking order (“V1-V2-V3-V4”, that reflects the expected canonical order of early visual areas) a post-hoc analysis was performed by manually examining ranking orders via a separate Chi-square test.

#### 2.4.4 Connectivity gradients within visual areas

Connectivity gradients within visual areas were calculated separately to define the functional organization within the visual cortex and how this organization differs in the blind group. Since the number of ROIs marked as “Visual” in the Glasser atlas is relatively low (26 unique regions), functional connectivity gradients were calculated vertex-wise. The vertices that correspond to “visual” areas according to Glasser areas were chosen in fsaverage5 mesh (Supplementary Figure 2). The selected vertices correspond to 2527 vertices out of 20484 that are available on fsaverage5 mesh. For each participant, preprocessed time series from those vertices were extracted and correlated with each other, resulting in 2527 × 2527 correlation matrices. The matrices were later normalized via Fisher’s Z transformation.

The mean correlation matrix of the sighted group was used to obtain the reference gradients that were used to align the individual gradients. The reference gradient and the individual gradients were calculated using the same method and parameters that were used to calculate whole-brain connectivity gradients (see Section 2.3). After calculating the gradient scores for each vertex, the age and sex effects are removed using a linear regression model, and the residuals are compared between sighted and blind groups via a two-sample t-test.

To assess the functional similarity of early visual regions, we compared the minimum functional distance between regions and evaluated group differences in these distances. The closer a region is to other regions, the higher functional similarity it represents. We used R^49^ and afex^50^ to perform a linear mixed effects analysis on the minimum distances. As fixed effects we entered group (blind vs sighted) and ROI (V1, V2, V3, V4), as well as their interaction terms into the linear mixed model (LMM). As random effects, we had intercepts for participants. P-values for fixed effects were obtained with the Satterthwaite method and were considered significant at p < .05. Post-hoc comparisons of significant interactions were conducted using approximate z-tests on the estimated marginal means (EMMs) using the emmeans package^51^. The resulting p-values were corrected for multiple comparisons following the procedure proposed by Holm (1979). Preliminary analyses of minimum distances and visual inspection of q-q plots of the residuals indicated deviations from normality. Thus, minimum distances were log transformed (log(minimum distance)) which led to a roughly normal distribution of the residuals.

#### 2.4.5 Structure-function coupling

For each participant, mean cortical thickness, curvature, sulcal depth, surface area, and volume of each 360 ROIs were extracted. To decrease the dimension of the data and account for all these structural measures at once, a principal component analysis (PCA) was applied to these five measures, and the first component was saved. This step created one structural component per participant.

Inter-subject correlation matrix on the structural component showed distinct similarity patterns across the three datasets, indicating a strong effect of the scanning site on the data. A one-way ANOVA test revealed that mean structural component values across 3 sites are statistically different (F(42,1)=20.36, p < .001; The test only included congenitally blind individuals, due to the absence of control subjects in Dataset 3). Therefore, the ComBat method, which is specifically designed for adjusting the data for the effect of scanning sites^52–54^ was used on the structural component through a publicly available Matlab toolbox (https://github.com/Jfortin1/ComBatHarmonization). Since Dataset 3 consists of congenitally blind participants only, it was not included in this step as data harmonization across sites requires balanced samples from two groups. After removing the participants from Dataset 3, 44 sighted and 27 congenitally participants remained in the pooled dataset. In addition to the site effects, the structural component was also corrected for age and sex during the same step.

A measure of structure-function coupling was calculated using Spearman’s rho rank correlation between the structural component and the gradient values for each ROI across sighted and blind participants. The structural component and the gradient values were previously corrected for age and sex. These resulted in two different structure-function coupling maps, for the sighted and blind groups respectively. For each group *i* and ROI *i*, the significance of the coupling measure *ρ_i_^j^* was obtained by Fischer-Z transforming the values,

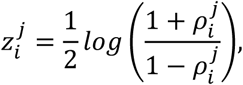

which are distributed according to a normal distribution. From the probability convolution theorem, it follows that the standardized structure-function coupling difference,

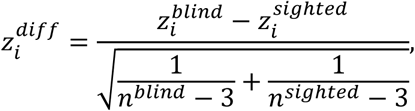

also follows a standard Gaussian distribution. *n^blind^* and *n^sighted^* are the number of subjects in each group. Then, the two-sided p-value can be easily calculated and FDR corrected using common statistical procedures. In addition to the ROI-wise comparison of the coupling values, coupling strength across the whole cortex was calculated. We defined the coupling strength as the mean absolute coupling value across the whole cortex. Mean absolute coupling values across gradients and both groups were compared via a two-way ANOVA.

In addition to the whole-brain ROI-based calculation of structure-function coupling, vertex-wise structure-function coupling within visual areas was calculated. The aim was to complement the vertex-wise gradient analysis within visual areas. First, PCA was applied to the same five structural measures (vertex-wise cortical thickness, curvature, surface area, volume, and sulcal depth) to create a common structural component (see above). The effect of the scanning site was adjusted for at this step using the Combat method^52^, and age and sex were included as covariates of no interest. Since Combat requires balanced samples from each site, Dataset 3 was removed from the analysis because it consisted of blind participants only. Each participant’s gradient value (after regressing out the effects of age and sex via linear regression) and the structural component were correlated with Spearman’s rank correlation within V1, V2, V3, and V4. At the end of this step, each participant had a correlation value for each vertex. Mean correlation values were calculated for each ROI. The mean correlation values were compared between sighted and blind groups for each ROI and gradient following the same procedure described in above.

## 3 Results

### 3.1 Demographics

The groups did not significantly differ in age (number of congenitally blind participants = 41, mean age = 36.71 ± 14.46, number of sighted participants = 44, mean age = 36.73 ± 13.79; t(83) = 0.004, p = .996) or sex distribution (blind = 21 male/20 female, sighted = 20 male/24 female; X^2^ = 2.28, p = .595).

### 3.2 Connectivity gradients across neocortex

Functional connectivity gradients revealed a canonical functional similarity profile of the cortical regions. The principal gradient (G1) showed that brain activity is maximally organized between the unimodal/transmodal cortices. The second axis (G2) showed the visual and sensorimotor/auditory regions, and the third axis (G3) showed the axis between the frontoparietal control network and the rest of the brain (Figure 1; see also, Supplementary Figure 1). On average, the first three gradients explained 59.68 % of the total variance for both groups (32.01% ± 0.07, 17.23% ± 0.03, and 10.45% ± 0.02, respectively). There were no differences in the explained variance of the gradients between the two groups (explained variances for each gradient were compared via a two-sample t-test, p > 0.05 for three gradients, corrected with FDR).

**Figure 1:**
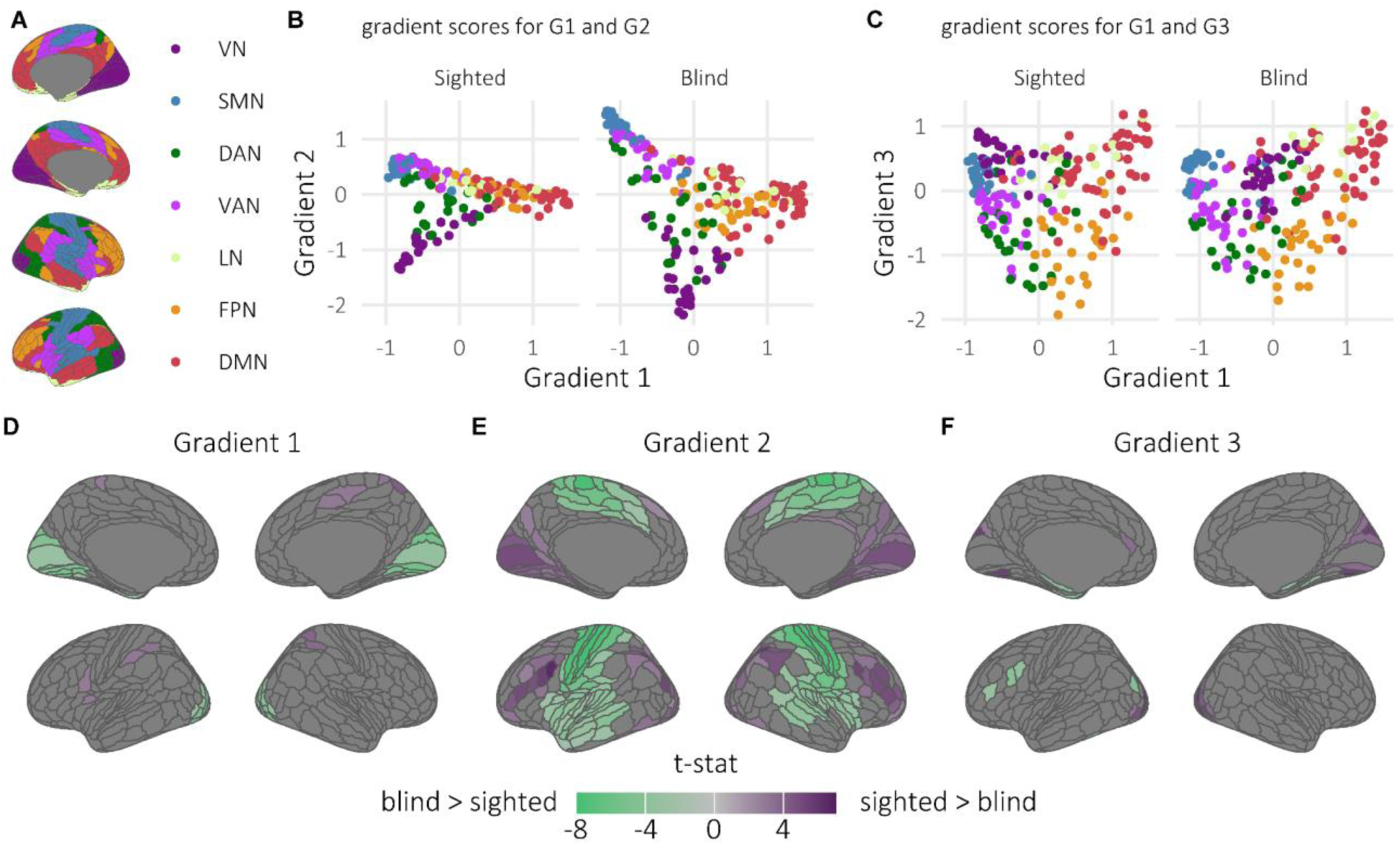
Differences in macroscale functional organization between blind and sighted individuals. **A** Projection of 7 functional networks defined by Yeo et al.^48^ on ROIs defined by Glasser et al.^44^ **B - C** Distribution of gradient values across G1 and G2 and across G1 and G3 in the groups of blind and sighted individuals. **D-E-F**: Results of the region-wise comparison of the gradient scores between groups of sighted and blind individuals (results were corrected for multiple comparisons with FDR at p < .05). VN = visual network; SMN = somatosensory/motor network; DAN = dorsal attention network; VAN = ventral attention network; LIN = limbic network; FPN = frontoparietal network; DMN = default mode network.

Region-wise comparisons of the gradient scores showed that in all three gradients, the groups of blind and sighted individuals significantly differed within visual regions (Figure 1D-F). In G1, the group of blind individuals had higher gradient scores in predominantly early, ventral, and lateral visual areas, suggesting that those areas are less segregated from transmodal areas in blind individuals (Table S1 and S4). In addition, we observed lower gradient scores for G1 in several regions of the somatosensory/motor and dorsal-attention networks in blind individuals, suggesting that the sensory-motor and the dorsal-attention networks are more segregated from the default-mode network in blind individuals than in sighted controls. In G2, blind individuals had lower gradient scores in predominantly early and ventral visual areas and higher gradient scores in the somatosensory/motor network, suggesting that in blind individuals early and ventral visual regions are more segregated from the somatosensory/motor network than in sighted controls (Table S2 and S4). In G3, blind individuals had lower scores in predominantly early, dorsal, and lateral visual areas than sighted controls, suggesting that these regions are closer to the frontoparietal network in blind individuals (Table S3 and S4).

Overall, the functional profile of early, ventral, and lateral visual areas of blind individuals becomes more similar to that of higher-order regions, and the distinction between visual areas and sensorimotor areas is increased. Moreover, early, dorsal and lateral visual areas of blind individuals seem to become more functionally similar to the frontoparietal network. In addition, sensorimotor areas shifted away from the origin of the gradient axis in G2 in blind individuals (Figure 1B), similar to what has been observed for visual regions.

To summarise the findings from the ROI analysis, we applied a well-established functional community decomposition (see Methods)^48^. Then we conducted a network-based z-score analysis of gradient scores for each of the first three gradients. This analysis revealed that in G1, blind individuals had higher gradient scores (i.e. less negative) in the visual network than sighted controls (t(51.499) = 3.897, p < .001) and higher gradient scores (i.e. more positive) in the sensory-motor network (t(51.925) = −4.943, p = .001) and the ventral attention network (t(51.116) = 2.693, p = .022, Figure 2A, see Table S5 in supplementary material for parameter estimates and confidence intervals). In G2, the group of blind individuals had lower gradient scores in the visual network (t(49.015) = 5.446, p< 0.001, Figure 2B) and the frontoparietal network (t(39.347) = 4.759, p < .001) than the group of sighted controls and higher gradient scores in the somatosensory/motor network (t(42.367) = −12.791, p < .001) and the ventral attention network (t(36.324) = −4.889, p < .001). Moreover, in G3 we observed that the group of blind individuals had lower gradient scores in the somatosensory/motor network (t(52.754) = 3.999, p < .001, Figure 2C) than sighted controls.

**Figure 2:**
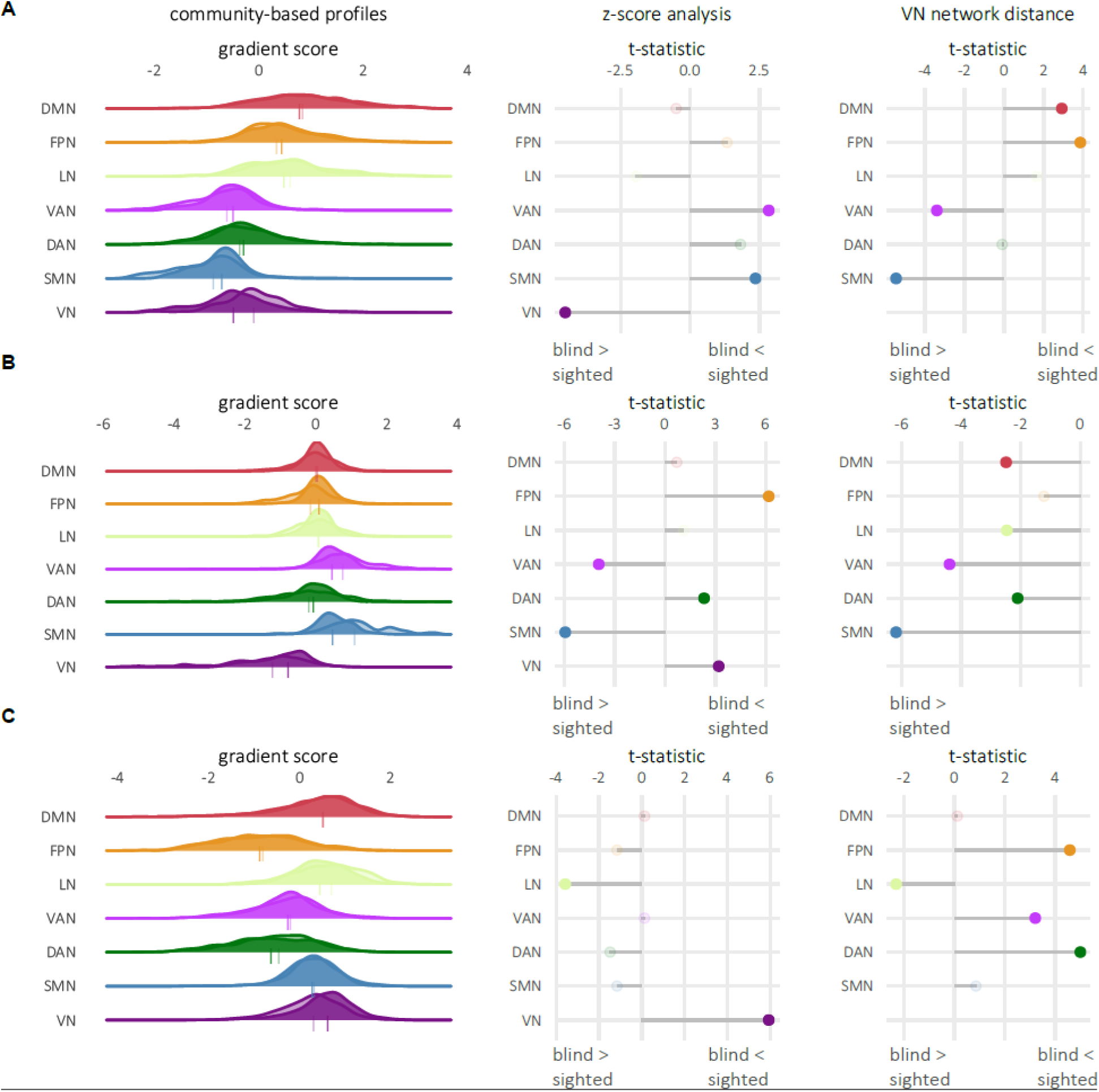
Differences in community based profiles between the groups of blind and sighted individuals. **A-B-C left panel**: Distribution of the gradient scores for each of the 7 functional networks (transparent distribution: group of blind individuals, non-transparent distribution: group of sighted controls). The tick below the density plots represents the mean gradient score. **A-B-C middle panel**: Group differences in the mean gradient scores across 7 functional networks. Significant differences are shown with full color, whereas non-significant differences are shown with transparent color. **A-B-C right panel**: Group differences in the distance between the visual and other functional networks. Significant differences are shown with full color, whereas non-significant differences are shown with transparent color. VN = visual network; SMN = somatosensory/motor network; DAN = dorsal attention network; ventral attention network = ventral attention network; LIN = limbic network; FPN = frontoparietal network; DMN = default mode network.

In the next step, we tested for potential group differences in the distance of the visual network and the rest of the networks. In G1, the visual network of blind individuals showed a greater distance to the somatosensory/motor network and the ventral attention network compared to sighted controls (group x network interaction: F(6, 498.00) = 19.24, p < .001, Figure 2A, see Table S6 and S7 for results of the post-hoc tests and EMMs), whereas the frontoparietal and default mode network were closer to the visual network in blind individuals than in sighted controls. In G2, the visual network of blind individuals was further apart from the somatosensory/motor, dorsal attention, ventral attention, frontoparietal, limbic, and the default mode network than the visual network of sighted controls (group x network interaction: F(6, 498.00) = 17.27, p < .001, Figure 2B, see Table S8 and S9 for results of the post-hoc tests and EMMs). In G3, the visual network of blind individuals was closer to the dorsal attention, ventral attention, and frontoparietal network than the visual network of sighted controls (group x network interaction: F(6, 498.00) = 11.23, p < .001, Figure 2C, see Table S10 and S11 for results of the post-hoc tests and EMMs), whereas the visual network of blind individuals showed greater distance to the limbic network than the visual network of sighted individuals. Overall, in G1 and G3, the visual network moved towards high-order cortices, while in G2 the visual network appeared more segregated.

The gradient range of G2 (defined as the maximum value of G2 minus the minimum value of G2) was higher in blind individuals (t(83) = 4.81, p < .001). No differences in gradient range between the groups were observed in G1 and G3 (all p > .05). Although the gradient range was related to the explained variance for both groups (Spearman’s Rho between gradient range and explained variance for G1: 0.85, G2: 0.43, G3: 0.26), the explained variances did not significantly differ between the groups of sighted and blind individuals (all ps > .05). Another variable correlated with gradient range for both groups was gradient variance (correlation between G1 range and variance: 0.96, G2: 0.93, G3: 0.91). Here, we observed that the variance of G2 was higher in blind individuals compared to sighted controls (t(83): 4.21, p < .001).

### 3.3 Hierarchical order of the early visual areas in the gradient space

Functional gradients reflect the hierarchical organization of cortical processing. Accordingly, gradient scores can be interpreted as indices of hierarchical ordering across cortical regions. Previous studies in enucleated macaques have shown that visual experience shapes the processing hierarchy within visual hierarchies^55^. To investigate whether a similar effect occurs in congenitally blind humans, we used gradient scores as a proxy for hierarchical ordering.

In sighted controls, the median ranking of the early visual cortex (EVC) in G1 showed the canonical “V1-V2-V3-V4” order 4265 times out of 10000 bootstrapped samples. This linear ranking order from V1 to V4 was not as frequent in the blind group (514 times out of 10000, Figure 3B, Table S12). For G2, both groups showed the “V1-V2-V3-V4” order as the most common one (6147 times for sighted controls, 5540 times for blind individuals, Figure 3B, Table S12). Lastly, for G3, the sighted group showed a relatively uniform distribution, having “V4-V3-V2-V1” order as the most common one (2334 times), whereas the blind group showed a dominant distribution for “V2-V1-V3-V4” order 6942 times (Figure 3B, Table S12). The complete details of the distribution are available in Table S12. These distributions imply that a linear order in the gradient space of G1 from V1 to V4 is available in sighted individuals, but not in blind individuals, while the canonical order remains largely unaltered in the G2 axis.

**Figure 3:**
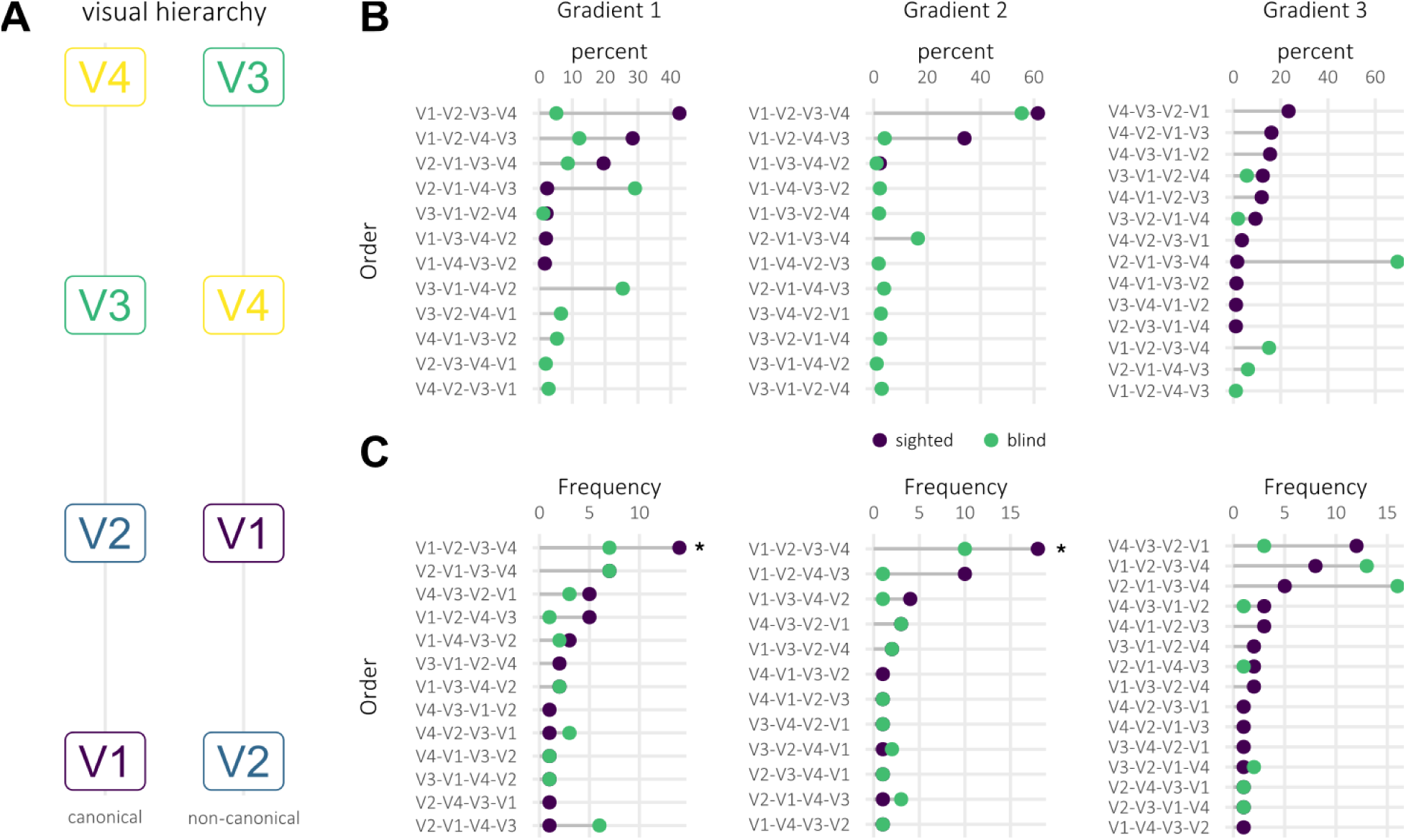
Differences in hierarchical organisation of the early visual cortex between congenitally blind individuals and sighted controls. **A** Hypothetical canonical and non-canonical hierarchy of the early visual cortex. **B** Percent distribution of the ranking orders across groups and gradients after bootstrapping. The bootstrapped samples showed “V1-V2-V3-V4” ranking order within early visual areas for the first two gradients. Only ranking orders with percent > 1 are displayed. **C** Frequency distribution of the ranking orders without bootstrapping. The canonical “V1-V2-V3-V4” distribution showed significant group differences for both G1 and G2 (p < .05, marked with asterisk).

In addition to bootstrapping, which enables us to inspect the most common ranking order estimate by resampling the dataset, we also directly compared the distributions and frequencies of different orders across the two groups without bootstrapping. Subject-wise rank orders showed different distributions across sighted and blind groups for all three gradients (G1: X^2^ = 19.01 p < .001; G2: X^2^ = 22.41 p < .001; G3 X^2^= 27.69 p= < .001). In G1, post-hoc comparison showed that the frequency of the canonical order (“V1-V2-V3-V4”) is different between the two groups (occurrence in the sighted group: 14 (31.81 % of whole sighted group), occurrence in the blind group: 7 ( 17.07% of the whole blind group; X^2^ = 17.19, p < .001). This finding is similar for G2 (occurrence in the sighted group: 18; 40.91% of the whole sighted sample, occurrence in the blind group: 10; 24.39% of the whole sample;X^2^ = 24.14, p < .001). In the blind group, the occurrence of the linear order in G2 at the subject-level shows a different pattern than that of sample-level (results of the bootstrapping). In G2, bootstrapping results indicate an intact canonical order, whereas the subject-level distribution shows that this order may not be as common in blind as in sighted individuals. Nonetheless, this reduction in occurrence could be attributed to a larger variability of the blind group rather than an inherent alteration of the hierarchical processing. Lastly, for G3, consistent with the bootstrapping results, the most common ranking order for the sighted group was “V4-V3-V2-V1” (12 times), and for the blind group “V2-V1-V3-V4” (16 times). A complete list of the distributions is available in Table S13.

### 3.4 Connectivity gradients within visual areas

Resting-state functional connectivity gradients can be used for parcellations of cortical functional areas. These parcellations have been shown to exhibit strong correspondence with task activation maps and with probabilistic V1 and V2^32,33^. Here, we apply this approach to assess whether functional arealization in the early visual cortex is shaped by visual experience.

The first three connectivity gradients within visual areas explained 49% of the variance in sighted controls (27.68% ± 10.33, 11.64% ± 2.52, 7.67% ± 1.94 for G1, G2, and G3 respectively). G1 did not show any difference between blind individuals and sighted controls (explained variance: 27.11 8.28, t(83): −0.31, p = .756), but G2 (explained variance: 15.14 ± 3.73, t(83): 5.16, p = .001) and G3 (explained variance: 8.94 ± 2.01, t(83): 2.91, p = .004) showed a higher explained variance in blind relative to sighted individuals.

In G1, the connectivity within visual areas is mainly organized between V1 and lateral occipital areas in sighted individuals. Along this axis, blind individuals showed lower gradient scores in V1, specifically in the occipital pole, and higher values in MT+ (Figure 4A), suggesting altered functional areal specialization in the group of blind individuals. G2 is organized along the posterior-anterior axis. In G2, the group of blind individuals had lower values in the dorsal stream and MT+. Lastly, G3 shows an organization between MT+ and the dorsal stream. Scores of the blind group increased in the dorsal stream and decreased in V1 (Figure 4A). The decomposition of the gradient scores by regions can be found in Supplementary Figure 3. Apart from the vertex-wise gradient scores, the groups of blind and sighted individuals also differed in gradient range and variance in gradient scores in G2, but not in G1 and G3.

**Figure 4:**
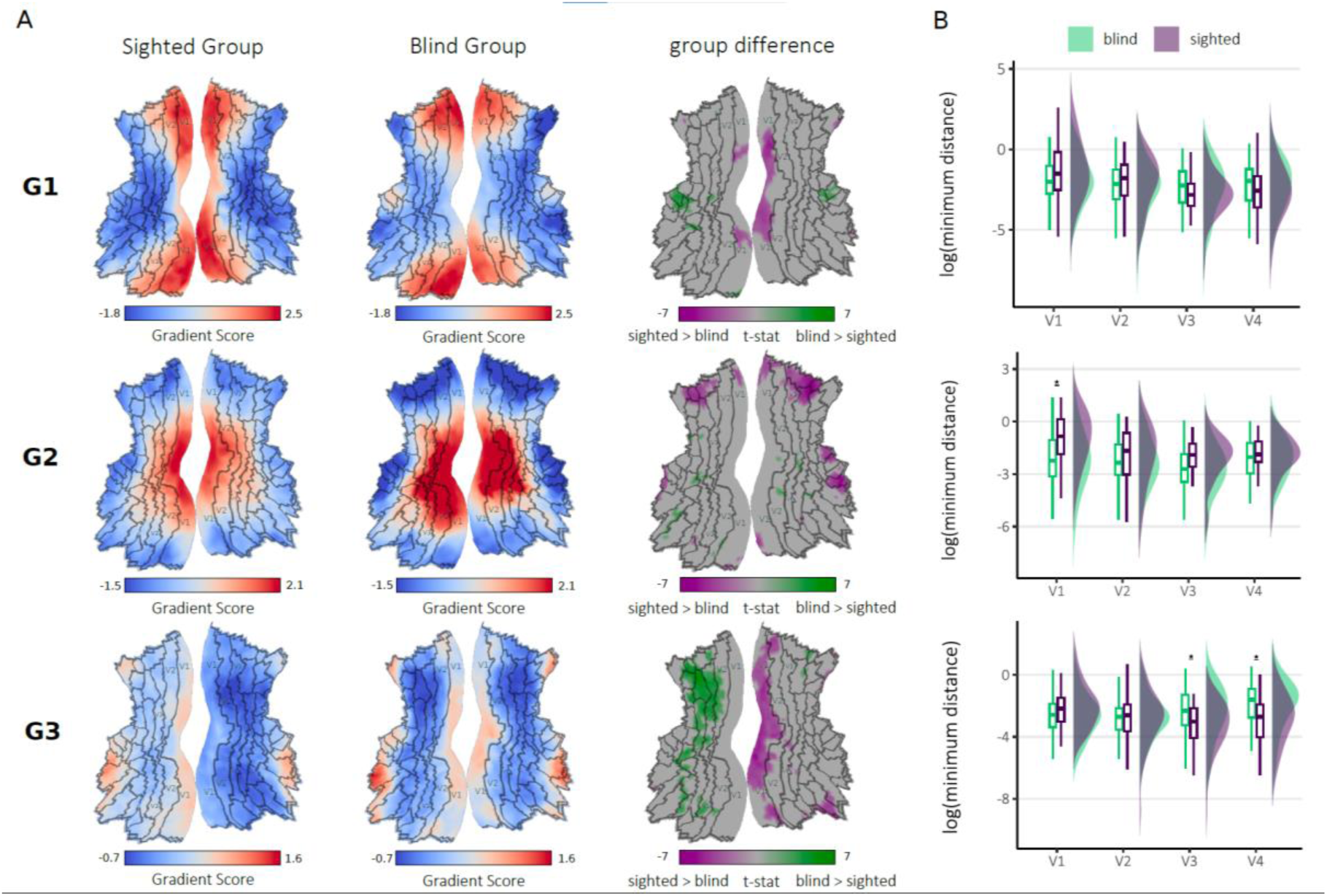
**A Functional organization within visual areas**. G1 shows that functional organization in visual areas is mainly defined by the functions from V1/V2 to the rest of the visual areas. G2 is shaped around the anterior-posterior axis, and G3 is organized within the dorsal stream. **B** Minimum functional distance to the rest of the early visual regions.

To summarise the findings from the vertex-wise analysis, we additionally assessed the minimal functional distance of each early visual area (V1, V2, V3, V4) to the other early visual areas. No significant group differences in log(minimum distance) were observed in G1 (Figure 4B, Table S14 and Table S15). In G2, the group of sighted individuals had a greater log(minimum distance) for V1 than the group of blind individuals (group x ROI interaction: (F(3, 249.00) = 4.52, p = .004, see Table S16 for the EMMs (non-transformed data for ease of interpretability) and Table S17 for the post-hoc tests (performed on the log scale), and Figure 4B). This suggests that in G2, V1 is more functionally distinct from the other areas of the EVC (V2, V3, V4) in sighted controls than blind individuals. In G3, the group of blind individuals had a greater log(minimum distance) for V3 and V4 than the group of sighted individuals (group x ROI interaction: (F(3, 249.00) = 9.66, p < .001, see Table S18 for the EMMs and Table S19 for the post-hoc tests). This suggests that in G3, V3 and V4 are more functionally distinct from V1 and V2 in blind individuals than in sighted controls.

### 3.5 Structure-function coupling

To study the coupling between structure and function in blind individuals, we obtained a low-dimensional representation of several structural measures (see Methods). The principal structural component created for each participant using their cortical thickness, volume, surface area, sulcal depth, and curvature explained 98.43% ± 0.41 of variance in both groups. Region-wise comparison between the structural component scores of the groups of sighted controls and blind individuals did not show any significant differences after correction for multiple comparisons. However, non-corrected p-values indicated that the group of blind individuals had lower structural scores in visual and sensorimotor areas, and higher scores in higher-order areas (Supplementary Figure 5A-B).

To assess structure–function coupling, we evaluated the relationship between the structural composite measure and the gradient scores from G1, G2, and G3 separately. The coupling between G1 and the structural component resembled the pattern of G1 itself, indicating a strong underlying structural mechanism for functional organization. Mean absolute coupling values across gradients and both groups were compared via a two-way ANOVA. The ANOVA model revealed a significant interaction between group and gradient (F(2,2154) = 63.62, p < .001, Table S20). Tukey-kramer post-hoc tests revealed that G1’s mean absolute coupling value was significantly higher in sighted controls than in blind individuals (mean absolute coupling value of sighted group = 0.31 ± 0.16, of blind group = 0.19 ± 0.13, estimated value of difference (EVD) = 0.11, p < .001, Figure 5C). A complete list of EVDs can be found in Table S21. When the coupling values for G1 in each ROI were compared between groups, visual (Figure 5B, regions colored with purple) and middle temporal areas (Figure 5B, regions colored in green, p < .05, corrected with FDR across whole-brain) showed a stronger structure-function coupling in the group of sighted controls (Table S22).

**Figure 5:**
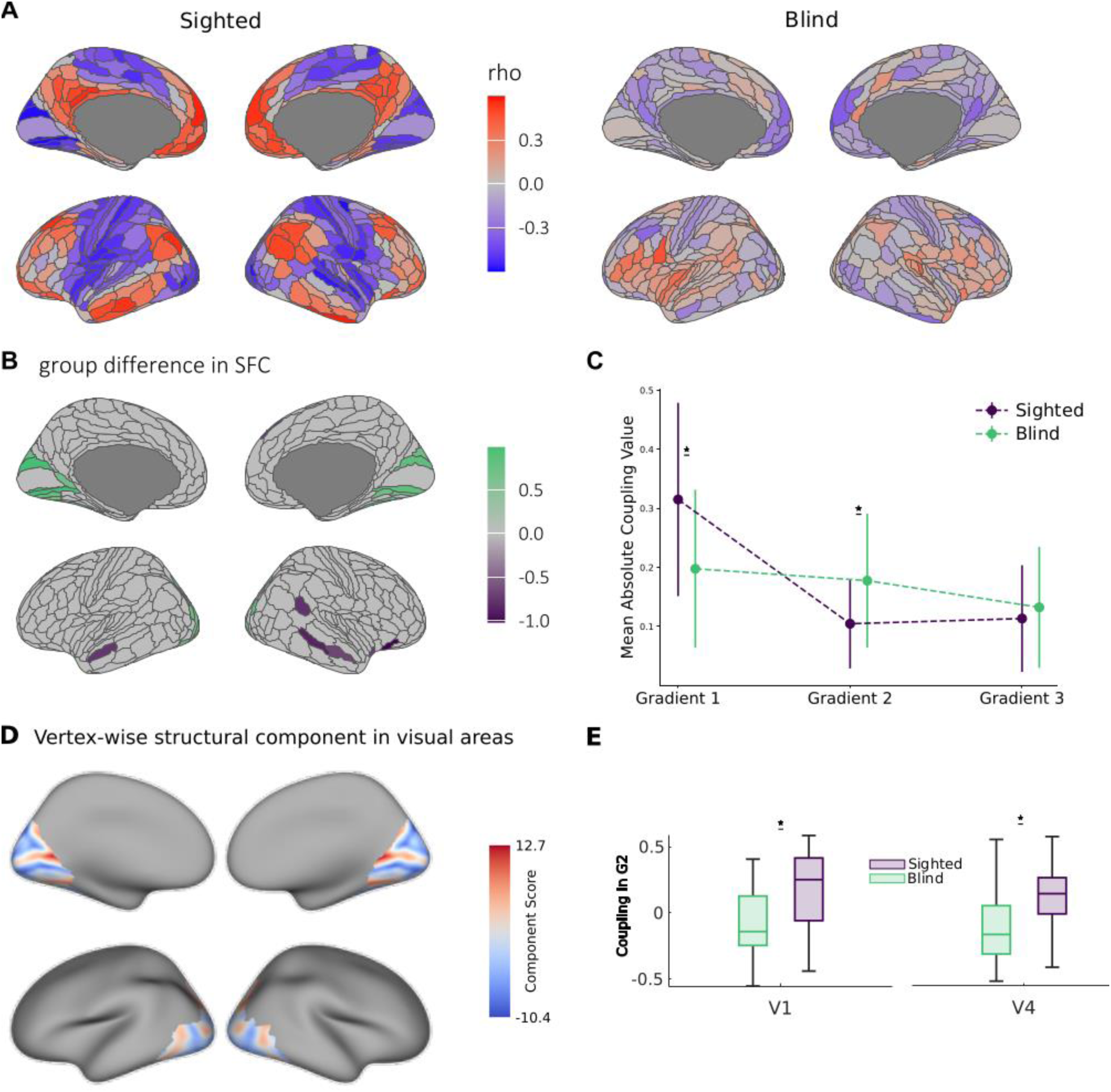
Correlation between G1 values and the structural components across groups. **A** Structure-function coupling values for groups of sighted controls and blind individuals. **B** Decreased structure-function coupling for G1 in the group of blind individuals in visual and temporal areas. **C** Comparison of the mean absolute coupling value across gradients and groups. The mean absolute coupling value derived from the coupling maps indicates the strength of the coupling across the cortex. The group of blind individuals shows weaker coupling with G1 and stronger coupling with G2. **D** Vertex-wise structural component within visual areas. **E** Correlation between the structural component shown in Panel D and mean coupling with G2 in V1 and V4 across two groups. The group of blind individuals showed less structure-function coupling compared to the group of sighted controls.

Tukey-Kramer post-hoc test following the ANOVA model above revealed that in sighted controls, the coupling between G2 and the structural component showed a similar pattern to G2 itself, that is, a positive coupling in somatosensory/motor areas and a negative coupling in visual areas (mean absolute coupling value: 0.10 ± 0.07). However, compared to G1, the overall coupling strength decreased significantly, indicating a weaker coupling mechanism for G2 (EVD = 0.21, p < .001, Table S21). These results indicate considerable but weaker structure-function coupling in G2 for the sighted control group compared to G1. By contrast, for the blind group, the coupling strength did not significantly differ between G1 and G2 (mean absolute coupling value for G1: 0.19 ± 0.13, for G2: 0.17 ± 0.11, EVD = 0.02, p = .188, Table S21). A significant difference was observed in mean absolute structure-function coupling in G2 between blind individuals and sighted controls (mean absolute coupling value of sighted group = 0.10 ± 0.07, of blind group = 0.17 ± 0.11, EVD = −0.07, p = < .001, Table S21). However, ROI-wise comparisons of the structure-function coupling values for G2 between sighted and blind groups did not reveal any significant differences.

The mean absolute structure-function coupling value in G3 for the sighted group was 0.11 ± 0.09, and 0.13 ± 0.10 for the blind group. Compared to G1, the overall coupling strength decreased significantly, indicating a weaker coupling mechanism for G3 (EVD = 0.201, p < .001, Table S21). No significant differences between groups were observed in the mean absolute structure-function coupling in G3 (sighted group = 0.11 ± 0.09, blind group = 0.13 ± 0.10, EVD = −0.01, p = .247, Table S21), nor in the ROI-wise comparisons.

The structural component explains 94.56% ± 1.41 of the variance across vertex-wise cortical thickness, volume, sulcal depth, surface area, and curvature within the visual ROIs. Since it was calculated on the vertex level, unlike the ROI-based component, it shows finer structural details, such as sulci and gyri (Figure 5D). The coupling between the structural component and the gradient scores across V1, V2, V3, and V4 showed that V1 and V4 have a decreased coupling with G2 for the blind group (mean V1-G2 coupling in sighted group: 0.16, 0.3, blind group: −0.08 ± 0.26, t(69): 3.65, p < .001; mean V4-G2 coupling in sighted group: 0.13 ± 0.2, in blind group: −0.08 ± 0.24, t(69): 3.95, p < .001, Figure 5E, correlations of every ROI with every gradient can be found in Supplementary Figure 4).

## 4 Discussion

A fundamental principle of human brain organization is the hierarchical arrangement within and between cortical networks. This hierarchical organization guides the flow of information and allows higher-order areas to process transmodal information that is unrelated to immediate sensory input^14,56^. Here, we investigated the extent to which this organization is shaped by sensory experience during development by comparing functional connectivity gradients and their coupling to structural measures in congenitally blind and sighted individuals.

Our findings suggest that the brain’s broad macroscale organization develops largely independent of visual experience. Both groups showed a principal gradient (G1) that extended from unimodal to transmodal areas, a second gradient (G2) that extended from somatosensory to visual areas, and a third gradient (G3) that differentiated the fronto-parietal network from other cortical regions. However, within this broadly preserved architecture, key differences emerged. In blind individuals, the visual network was functionally more segregated from the sensorimotor network (G2) and more integrated with transmodal networks (G1) and the frontoparietal network (G3). These shifts may support the recruitment of visual cortices for higher-order cognitive functions, such as language^4,57–59^, memory^5^, and executive control^5^ in blind individuals.

These macroscale shifts were accompanied by alterations in the internal organization of the visual cortices. Because functional gradients capture the hierarchical organization of cortical processing, gradient scores can be interpreted as indices of hierarchical ordering across regions. To test whether visual experience shapes cortical hierarchies, we derived a measure of hierarchical ordering from the relative positions of early visual areas (V1–V4) along the first three gradients and compared these rankings between the sighted and blind groups. In the sighted group, the first gradient (G1) showed a very consistent linear order from V1 to V4, mirroring the canonical visual hierarchy. This ranking was much less consistent in the blind group. This variability may reflect increased interindividual heterogeneity and a reorganization of connectivity in the absence of feedforward visual input^60^. Notably, V1 was less distinct from V2, V3, and V4 in blind individuals. Previous studies in enucleated macaques have reported the development of a “hybrid cortex” between V1 and V2 that combines cytoarchitectonic characteristics of both areas^55,61^. It is possible that the reduced functional differentiation observed in blind participants reflects a similar reorganization.

Gradient-based analyses of functional organization within the visual cortex further revealed altered areal specialization in blind individuals. In sighted individuals, gradient maps closely matched probabilistic V1– V2 borders, whereas this correspondence was reduced in blind individuals. V1 appeared functionally subdivided in blind individuals, with the occipital pole clustering with ventral occipitotemporal areas and peripheral V1 clustering with dorsal visual regions. These findings suggest that visual experience is critical for refining areal borders and shaping cortical gradients, particularly within the early visual cortex. The occipital pole appears especially sensitive to sensory input during development, showing both decreased functional differentiation and structural changes, such as altered cortical thickness reported in previous studies^62^, while the calcarine sulcus remains largely unaffected. Together, these observations raise the possibility that the occipital pole in blind individuals may resemble the hybrid cortex described in enucleated macaques, although this remains speculative.

Next, we examined the coupling between structural and functional cortical organization. Structural organization was quantified using a composite measure of cortical thickness, surface area, curvature, volume, and sulcal depth. Functional organization was assessed based on gradient scores derived from the first three principal gradients (G1, G2, and G3). Across the cortex, G1 exhibited strong coupling with structural features, but this coupling was markedly reduced in visual and temporal regions of blind individuals. Previous studies have shown that structure–function coupling in visual areas emerges early in life and strengthens throughout development^63,64^. Our findings indicate that this developmental trajectory is shaped by sensory experience: in the absence of visual input, structure–function coupling is reduced, particularly in visual cortices. Increased myelination and a lower excitation–inhibition (E/I) ratio have been linked to tighter structure–function coupling^65^ and there is evidence to suggest that the E/I ratio is increased^66,67^ and myelination is reduced within early visual areas of congenitally blind individuals. Thus, the reduced structure-function coupling we observed in the group of blind individuals may be the result of reduced myelination and altered E/I balance.

It should be noted that the reduction in structure-function coupling that we observed in blind individuals may also be due to methodological factors. This entails the assumption that functional areas can be mapped to identical spatial locations in all participants^68^. However, our data suggest altered functional areal boundaries and hierarchical organization in the blind group, especially within the early visual cortex. Misalignment between structural and functional parcels may therefore obscure true coupling relationships. Addressing this challenge will require individualized parcellations derived from resting-state data that better reflect functional topography in sensory-deprived populations^32,34,69^.

## Conclusion

In summary, our results show that the broad macroscale hierarchical organization of the brain is largely preserved in the absence of vision, suggesting an innate macroscale hierarchy. However, distinct alterations in both global gradient patterns and functional connectivity within the visual cortex in blind individuals highlight the importance of visual experience in fine-tuning these cortical hierarchies and in refining the borders between functional regions. Our findings underscore a dynamic interplay between intrinsic developmental processes and sensory experience in shaping cortical networks, particularly within visual networks.

## Data and code availability

The scripts used in this study are available at https://github.com/KobaCemal/BlindnessGradients and https://github.com/LenaStroh/BlindnessGradients. Dataset 2 is freely available at https://doi.org/10.3886/E198832V1

## Acknowledgements

Research supported by the Polish National Science Centre (NCN) grant no: 2018/30/A/HS6/00595 to Marcin Szwed. Cemal Koba and Joan Falcó-Roget were supported by the project of the Minister of Science and Higher Education “Support for the activity of Centers of Excellence established in Poland under Horizon 2020” on the basis of the contract number MEiN/2023/DIR/379, and the European Union’s Horizon 2020 research and innovation programme under grant agreement No 857533, and by Sano project carried out within the International Research Agendas programme of the Foundation for Polish Science, co-financed by the European Union under the European Regional Development Fund. Cemal Koba and Joan Falcó-Roget gratefully acknowledge Polish high-performance computing infrastructure PLGrid (HPC Center: ACK Cyfronet AGH) for providing computer facilities and support within computational grant no. PLG/2024/017108. Olivier Collignon is a senior researcher at FRS-FNRS in Belgium. We would like to thank all of our participants without whom this research would not have been possible.

## Supplementary Materials

Supplementary tables can be found at: [Supplementary Tables](https://1drv.ms/w/s!Ak2VRTvRuUzxs-57qsZITA1KZmxlQQ?e=PzDruZ)

**Supplementary Figure 1:**
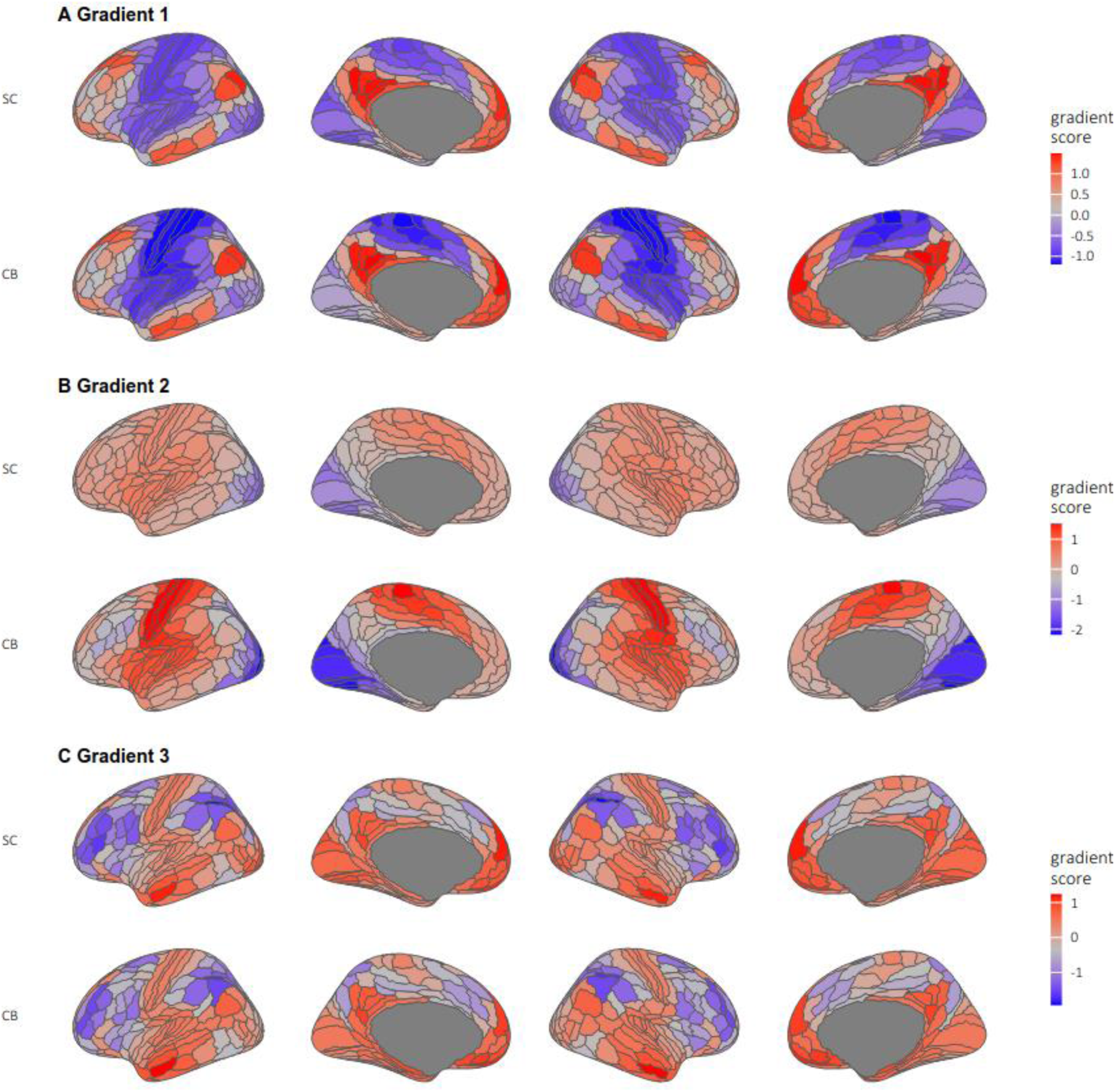
First three mean gradients in sighted and blind groups. SC: Sighted Control Group, CB: Congenitally Blind Group

**Supplementary Figure 2:**
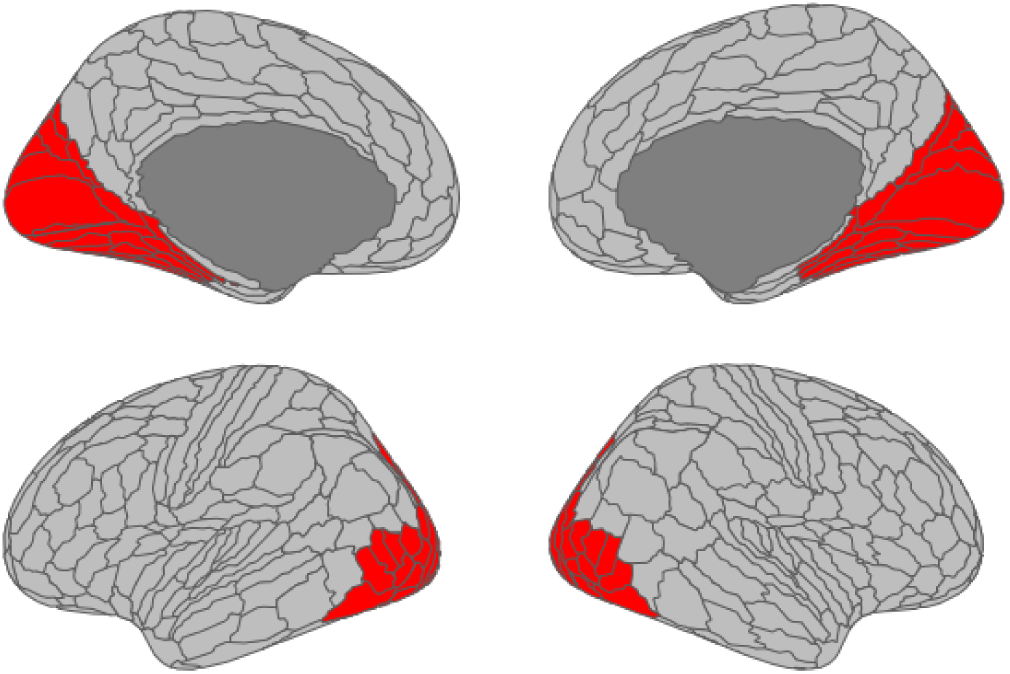
Network mask that is used to calculate connectivity gradients within visual areas.

**Supplementary Figure 3:**
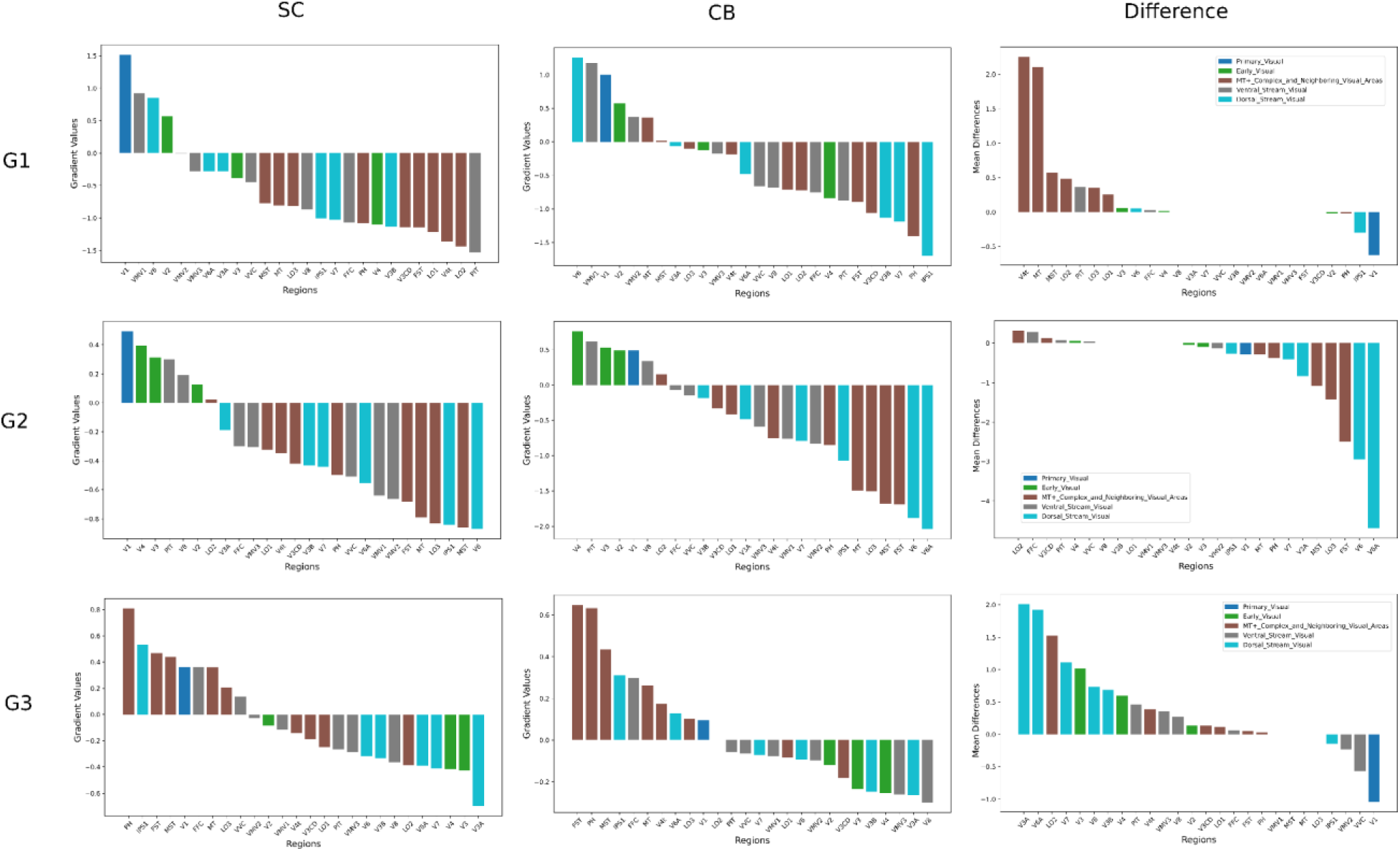
Decomposition of the vertex-wise gradients and their differences across visual regions. Left column: Mean gradients values across regions for sighted group. Mid panel: Mean gradients values across regions for blind group. Right column: Mean t-score of differences across regions (blind-sighted). SC: Sighted Control Group, CB: Congenitally Blind Group

**Supplementary Figure 4:**
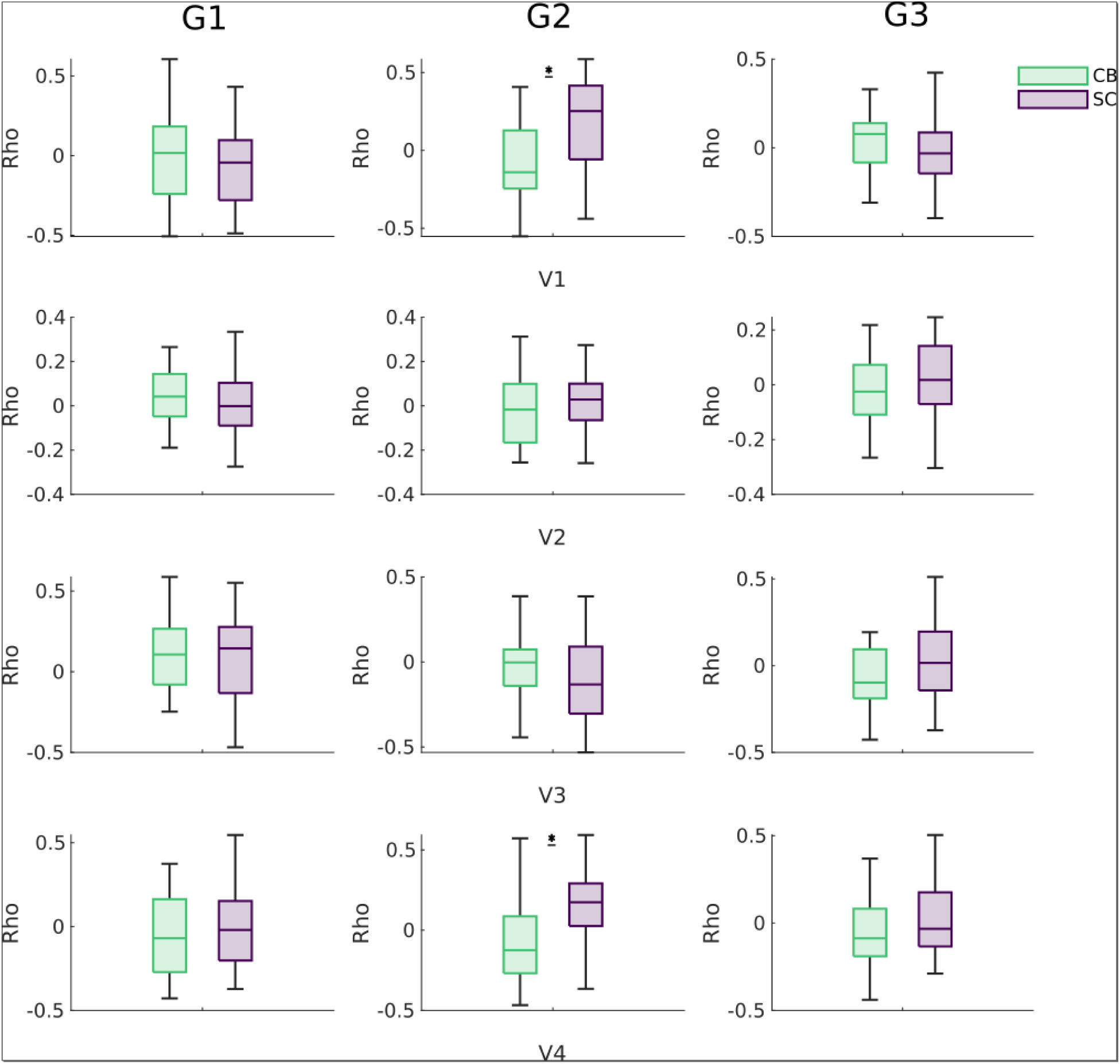
Structure-function coupling within visual areas across gradients. Asterisk indicates significant difference (p<0.05, corrected with FDR). SC: Sighted Control Group, CB: Congenitally Blind Group

**Supplementary Figure 5:**
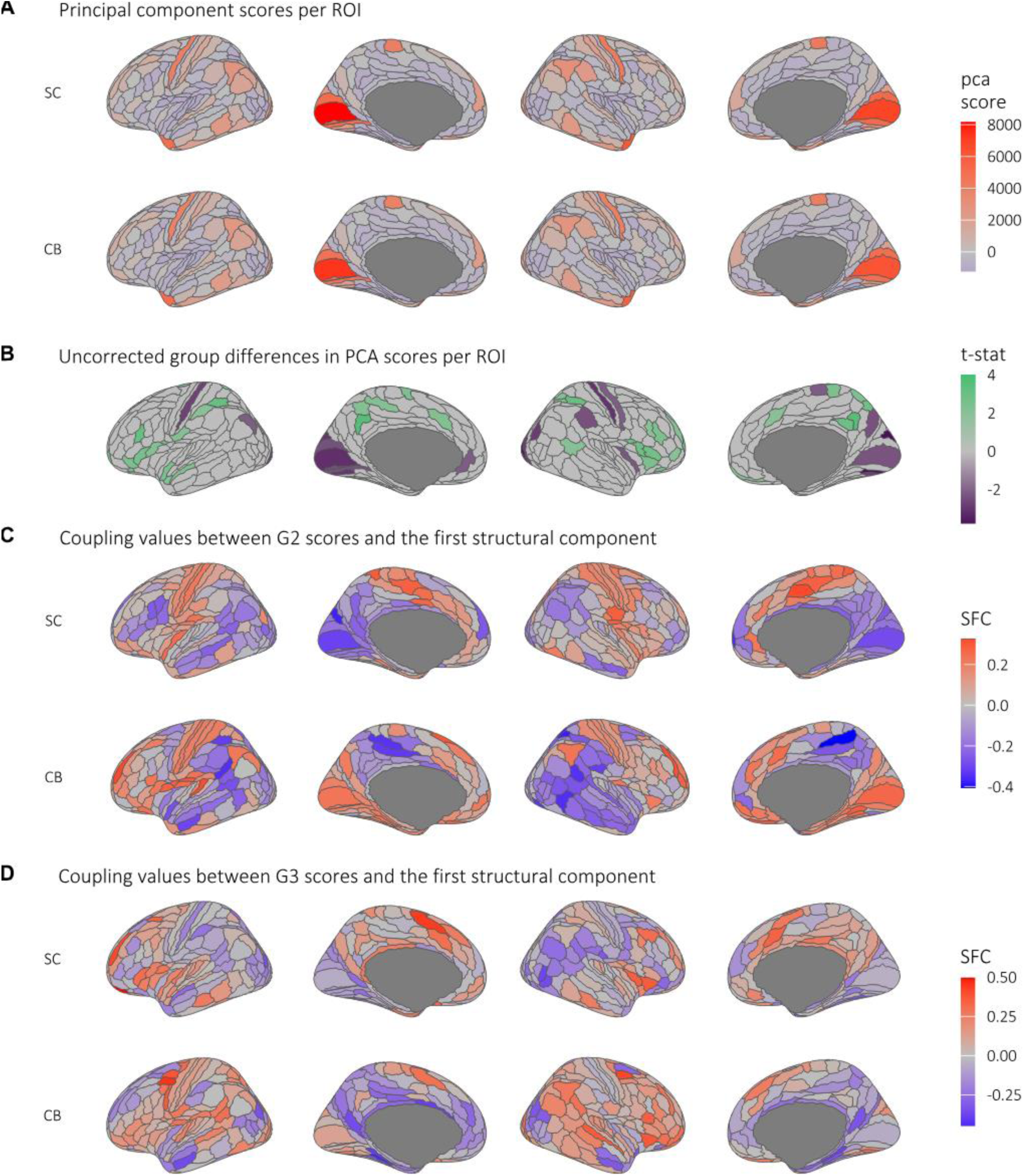
**A** Principal structural component for sighted and blind groups. **B** Uncorrected difference map of the structural component (blind>sighted). The group of blind individuals showed lower component scores in visual and sensorimotor areas, and higher scores in higher-order areas (p < .05) **C-D** Coupling values between the gradient scores of G2 and G3 and the first structural component. SC: Sighted Control Group, CB: Congenitally Blind Group

## References

1. Mesulam, M. M. From sensation to cognition. Brain 121 (Pt 6), 1013–1052 (1998).

2. Margulies, D. S. et al. Situating the default-mode network along a principal gradient of macroscale cortical organization. PNAS 113, 12574–12579 (2016).

3. Huntenburg, J. M., Bazin, P.-L. & Margulies, D. S. Large-Scale Gradients in Human Cortical Organization. Trends in Cognitive Sciences 22, 21–31 (2018).

4. Bedny, M., Pascual-Leone, A., Dodell-Feder, D., Fedorenko, E. & Saxe, R. Language processing in the occipital cortex of congenitally blind adults. Proceedings of the National Academy of Sciences 108, 4429–4434 (2011).

5. Abboud, S. & Cohen, L. Distinctive Interaction Between Cognitive Networks and the Visual Cortex in Early Blind Individuals. 18 (2019) doi:10.1093/cercor/bhz%20006.

6. Striem-Amit, E. et al. Functional connectivity of visual cortex in the blind follows retinotopic organization principles. Brain 138, 1679–1695 (2015).

7. Kanjlia, S., Pant, R. & Bedny, M. Sensitive Period for Cognitive Repurposing of Human Visual Cortex. http://biorxiv.org/lookup/doi/10.1101/402321 (2018) doi:10.1101/402321.

8. Liu, Y. et al. Whole brain functional connectivity in the early blind. Brain 130, 2085–2096 (2007).

9. Wang, D. et al. Altered resting-state network connectivity in congenital blind. Human Brain Mapping 35, 2573–2581 (2014).

10. Büchel, C. Cortical hierarchy turned on its head. Nat Neurosci 6, 657–658 (2003).

11. Hölig, C., Föcker, J., Best, A., Röder, B. & Büchel, C. Brain systems mediating voice identity processing in blind humans: Voice Identity Processing in Blind Humans. Hum. Brain Mapp. 35, 4607– 4619 (2014).

12. Wang, X. et al. Domain Selectivity in the Parahippocampal Gyrus Is Predicted by the Same Structural Connectivity Patterns in Blind and Sighted Individuals. J. Neurosci. 37, 4705–4716 (2017).

13. Dormal, G., Rezk, M., Yakobov, E., Lepore, F. & Collignon, O. Auditory motion in the sighted and blind: Early visual deprivation triggers a large-scale imbalance between auditory and “visual” brain regions. NeuroImage 134, 630–644 (2016).

14. Margulies, D. S. et al. Situating the default-mode network along a principal gradient of macroscale cortical organization. Proc. Natl. Acad. Sci. U.S.A. 113, 12574–12579 (2016).

15. Markov, N. T. & Kennedy, H. The importance of being hierarchical. Current Opinion in Neurobiology 23, 187–194 (2013).

16. Mesulam, M. From sensation to cognition. Brain 121, 1013–1052 (1998).

17. Taylor, H. P. et al. Functional Hierarchy of the Human Neocortex from Cradle to Grave. Preprint at 10.1101/2024.06.14.599109 (2024).

18. Zhang, J. et al. Intrinsic Functional Connectivity is Organized as Three Interdependent Gradients. Sci Rep 9, 15976 (2019).

19. Baum, G. L. et al. Development of structure–function coupling in human brain networks during youth. Proc Natl Acad Sci USA 117, 771–778 (2020).

20. Paquola, C. et al. Microstructural and functional gradients are increasingly dissociated in transmodal cortices. PLOS Biology 17, e3000284 (2019).

21. Preti, M. G. & Van De Ville, D. Decoupling of brain function from structure reveals regional behavioral specialization in humans. Nat Commun 10, 4747 (2019).

22. Vázquez-Rodríguez, B. et al. Gradients of structure–function tethering across neocortex. Proceedings of the National Academy of Sciences 116, 21219–21227 (2019).

23. Czarnecka, M. et al. Association between Cortical Thickness and Functional Response to Linguistic Processing in the Occipital Cortex of Early Blind Individuals. 2025.04.17.645592 Preprint at 10.1101/2025.04.17.645592 (2025).

24. Anurova, I., Renier, L. A., De Volder, A. G., Carlson, S. & Rauschecker, J. P. Relationship Between Cortical Thickness and Functional Activation in the Early Blind. Cereb. Cortex 25, 2035–2048 (2015).

25. Haak, K. V., Marquand, A. F. & Beckmann, C. F. Connectopic mapping with resting-state fMRI. NeuroImage 170, 83–94 (2018).

26. Watson, D. M. & Andrews, T. J. An evaluation of how connectopic mapping reveals visual field maps in V1. Sci Rep 12, 16249 (2022).

27. Ngo, G. N., Haak, K. V., Beckmann, C. F. & Menon, R. S. Mesoscale hierarchical organization of primary somatosensory cortex captured by resting-state-fMRI in humans. NeuroImage 235, 118031 (2021).

28. Tian, Y. & Zalesky, A. Characterizing the functional connectivity diversity of the insula cortex: Subregions, diversity curves and behavior. NeuroImage 183, 716–733 (2018).

29. Wang, R. et al. Functional connectivity gradients of the insula to different cerebral systems. Human Brain Mapping 44, 790–800 (2023).

30. Shen, Y. et al. Functional connectivity gradients of the cingulate cortex. Commun Biol 6, 1–9 (2023).

31. Song, Y. et al. Functional hierarchy of the angular gyrus and its underlying genetic architecture. Human Brain Mapping 44, 2815–2828 (2023).

32. Gordon, E. M. et al. Generation and Evaluation of a Cortical Area Parcellation from Resting-State Correlations. Cereb. Cortex 26, 288–303 (2016).

33. Wig, G. S., Laumann, T. O. & Petersen, S. E. An approach for parcellating human cortical areas using resting-state correlations. NeuroImage 93, 276–291 (2014).

34. Pelland, M. et al. State-dependent modulation of functional connectivity in early blind individuals. NeuroImage 147, 532–541 (2017).

35. Gorgolewski, K. J. et al. The brain imaging data structure, a format for organizing and describing outputs of neuroimaging experiments. Sci Data 3, 160044 (2016).

36. Bedny, M. & Tian, M. Blindness Resting State. ICPSR - Interuniversity Consortium for Political and Social Research 10.3886/E198832V1 (2024).

37. Tian, M., Xiao, X., Hu, H., Cusack, R. & Bedny, M. Visual experience shapes functional connectivity between occipital and non-visual networks. eLife 13, (2024).

38. Cruces, R. R. et al. Micapipe: A pipeline for multimodal neuroimaging and connectome analysis. NeuroImage 263, 119612 (2022).

39. Cox, R. W. AFNI: software for analysis and visualization of functional magnetic resonance neuroimages. Comput Biomed Res 29, 162–173 (1996).

40. Jenkinson, M., Beckmann, C. F., Behrens, T. E. J., Woolrich, M. W. & Smith, S. M. FSL. Neuroimage 62, 782–790 (2012).

41. Avants, B. B., Epstein, C. L., Grossman, M. & Gee, J. C. Symmetric diffeomorphic image registration with cross-correlation: Evaluating automated labeling of elderly and neurodegenerative brain. Medical Image Analysis 12, 26–41 (2008).

42. Dale, A. M., Fischl, B. & Sereno, M. I. Cortical Surface-Based Analysis: I. Segmentation and Surface Reconstruction. NeuroImage 9, 179–194 (1999).

43. Henschel, L. et al. FastSurfer - A fast and accurate deep learning based neuroimaging pipeline. NeuroImage 219, 117012 (2020).

44. Glasser, M. F. et al. A multi-modal parcellation of human cerebral cortex. Nature 536, 171–178 (2016).

45. Satterthwaite, T. D. et al. An improved framework for confound regression and filtering for control of motion artifact in the preprocessing of resting-state functional connectivity data. Neuroimage 64, 240–256 (2013).

46. Vos de Wael, R., et al. BrainSpace: a toolbox for the analysis of macroscale gradients in neuroimaging and connectomics datasets. Commun Biol 3, 1–10 (2020).

47. Worsley, K. et al. SurfStat: A Matlab toolbox for the statistical analysis of univariate and multivariate surface and volumetric data using linear mixed effects models and random field theory. NeuroImage 47, S102 (2009).

48. Yeo, B. T. T. et al. The organization of the human cerebral cortex estimated by intrinsic functional connectivity. Journal of Neurophysiology 106, 1125–1165 (2011).

49. Posit team. Rstudio: Integrated. Posit Software, PBC (2024).

50. Singmann, H., Bolker, B., Westfall, J., Aust, F. & Ben-Shachar, M. S. _afex: Analysis of Factorial Experiments_. (2023).

51. Lenth, R. V. _emmeans: Estimated Marginal Means, aka Least-Squares Means_. (2023).

52. Fortin, J.-P. et al. Harmonization of multi-site diffusion tensor imaging data. NeuroImage 161, 149– 170 (2017).

53. Fortin, J.-P. et al. Harmonization of cortical thickness measurements across scanners and sites. Neuroimage 167, 104–120 (2018).

54. Johnson, W. E., Li, C. & Rabinovic, A. Adjusting batch effects in microarray expression data using empirical Bayes methods. Biostatistics 8, 118–127 (2007).

55. Magrou, L. et al. How Areal Specification Shapes the Local and Interareal Circuits in a Macaque Model of Congenital Blindness. Cerebral Cortex 28, 3017–3034 (2018).

56. Paquola, C. et al. The architecture of the human default mode network explored through cytoarchitecture, wiring and signal flow. Nat Neurosci 1–11 (2025) doi:10.1038/s41593-024-01868-0.

57. Dzięgiel-Fivet, G. et al. Neural network for Braille reading and the speech-reading convergence in the blind: Similarities and differences to visual reading. NeuroImage 231, 117851 (2021).

58. Kim, J. S., Kanjlia, S., Merabet, L. B. & Bedny, M. Development of the Visual Word Form Area Requires Visual Experience: Evidence from Blind Braille Readers. J. Neurosci. 37, 11495–11504 (2017).

59. Lane, C., Kanjlia, S., Omaki, A. & Bedny, M. ‘Visual’ Cortex of Congenitally Blind Adults Responds to Syntactic Movement. Journal of Neuroscience 35, 12859–12868 (2015).

60. Sen, S. et al. The Role of Visual Experience in Individual Differences of Brain Connectivity. J. Neurosci. 42, 5070–5084 (2022).

61. Rakic, P., Suner, I. & Williams, R. W. A novel cytoarchitectonic area induced experimentally within the primate visual cortex. Proceedings of the National Academy of Sciences 88, 2083–2087 (1991).

62. Stroh, A.-L. et al. Blind individuals’ enhanced ability to sense their own heartbeat is related to the thickness of their occipital cortex. Cerebral Cortex 34, bhae324 (2024).

63. Hong, Y. et al. Structural and functional connectome relationships in early childhood. Developmental Cognitive Neuroscience 64, 101314 (2023).

64. Natu, V. S. et al. Infants’ cortex undergoes microstructural growth coupled with myelination during development. Commun Biol 4, 1191 (2021).

65. Fotiadis, P. et al. Myelination and excitation-inhibition balance synergistically shape structure-function coupling across the human cortex. Nat Commun 14, 6115 (2023).

66. Coullon, G. S. L., Emir, U. E., Fine, I., Watkins, K. E. & Bridge, H. Neurochemical changes in the pericalcarine cortex in congenital blindness attributable to bilateral anophthalmia. J Neurophysiol 114, 1725–1733 (2015).

67. Rączy, K., et al. Typical Resting State Activity of the Brain Requires Visual Input during an Early Sensitive Period. http://biorxiv.org/lookup/doi/10.1101/2021.06.09.446724 (2021) doi:10.1101/2021.06.09.446724.

68. Suárez, L. E., Markello, R. D., Betzel, R. F. & Misic, B. Linking Structure and Function in Macroscale Brain Networks. Trends in Cognitive Sciences 24, 302–315 (2020).

69. Tu, J. C. et al. Early Life Neuroimaging: The Generalizability of Cortical Area Parcellations Across Development. Preprint at 10.1101/2024.09.09.612056 (2024).

