## Supplementary tables for "Visual experience shapes functional connectome gradients"

**Table S1 - Regions that show significant differences gradient 1 scores between CB-SC groups.**

| region | hemi | yeo | stream | estimate | parameter | statistic | p (uncorr) | p (corr) |
|---|---|---|---|---|---|---|---|---|
| V3A | left | VIS | dorsal | 0.447 | 80.351 | 3.660 | < 0.001 | 0.006 |
| V3A | right | VIS | dorsal | 0.537 | 79.675 | 4.314 | < 0.001 | 0.001 |
| V3B | left | VIS | dorsal | 0.656 | 78.468 | 4.841 | < 0.001 | < 0.001 |
| V3B | right | VIS | dorsal | 0.577 | 79.093 | 4.283 | < 0.001 | 0.001 |
| V7 | left | VIS | dorsal | 0.430 | 81.819 | 3.490 | 0.001 | 0.009 |
| V1 | left | VIS | evc | 0.416 | 81.057 | 3.241 | 0.002 | 0.018 |
| V1 | right | VIS | evc | 0.408 | 82.986 | 2.859 | 0.005 | 0.039 |
| V2 | left | VIS | evc | 0.721 | 78.664 | 5.523 | < 0.001 | < 0.001 |
| V2 | right | VIS | evc | 0.638 | 80.736 | 4.487 | < 0.001 | 0.001 |
| V3 | left | VIS | evc | 0.621 | 82.925 | 5.348 | < 0.001 | < 0.001 |
| V3 | right | VIS | evc | 0.627 | 82.602 | 5.166 | < 0.001 | < 0.001 |
| V4 | left | VIS | evc | 0.669 | 82.738 | 5.439 | < 0.001 | < 0.001 |
| V4 | right | VIS | evc | 0.645 | 82.446 | 5.071 | < 0.001 | < 0.001 |
| LO1 | left | VIS | lateral | 0.514 | 82.200 | 3.883 | < 0.001 | 0.003 |
| LO1 | right | VIS | lateral | 0.489 | 81.603 | 3.642 | < 0.001 | 0.006 |
| LO2 | left | VIS | lateral | 0.547 | 82.065 | 4.682 | < 0.001 | < 0.001 |
| LO2 | right | VIS | lateral | 0.62 | 82.489 | 4.809 | < 0.001 | < 0.001 |
| V3CD | left | VIS | lateral | 0.621 | 81.492 | 4.450 | < 0.001 | 0.001 |
| V3CD | right | VIS | lateral | 0.537 | 82.916 | 3.701 | < 0.001 | 0.005 |
| V4t | left | VIS | lateral | 0.400 | 82.245 | 3.144 | 0.002 | 0.022 |
| V4t | right | VIS | lateral | 0.422 | 79.910 | 2.995 | 0.004 | 0.032 |
| FFC | left | VIS | ventral | 0.580 | 81.860 | 4.406 | < 0.001 | 0.001 |
| FFC | right | VIS | ventral | 0.470 | 82.424 | 3.933 | < 0.001 | 0.003 |
| PIT | left | VIS | ventral | 0.623 | 82.700 | 5.150 | < 0.001 | < 0.001 |
| PIT | right | VIS | ventral | 0.633 | 82.754 | 5.095 | < 0.001 | < 0.001 |
| V8 | left | VIS | ventral | 0.607 | 82.845 | 5.021 | < 0.001 | < 0.001 |
| V8 | right | VIS | ventral | 0.581 | 81.759 | 4.479 | < 0.001 | 0.001 |
| VMV1 | left | VIS | ventral | 0.702 | 77.719 | 5.175 | < 0.001 | < 0.001 |
| VMV1 | right | VIS | ventral | 0.832 | 77.323 | 5.448 | < 0.001 | < 0.001 |
| VMV2 | right | VIS | ventral | 0.391 | 82.950 | 2.879 | 0.005 | 0.038 |
| VMV3 | left | VIS | ventral | 0.438 | 82.881 | 3.100 | 0.003 | 0.024 |
| VMV3 | right | VIS | ventral | 0.506 | 82.992 | 3.836 | < 0.001 | 0.003 |
| VVC | left | VIS | ventral | 0.510 | 82.954 | 3.951 | < 0.001 | 0.003 |
| VVC | right | VIS | ventral | 0.444 | 81.540 | 3.221 | 0.002 | 0.018 |
| 24dd | left | SMN | | -0.348 | 74.050 | -2.959 | 0.004 | 0.032 |
| 24dv | left | SMN | | -0.354 | 70.800 | -2.976 | 0.004 | 0.032 |
| 5L | left | SMN | | -0.337 | 80.121 | -3.129 | 0.002 | 0.023 |
| 5m | right | SMN | | -0.344 | 82.984 | -2.895 | 0.005 | 0.037 |
| 7AL | left | DAN | | -0.306 | 73.966 | -3.261 | 0.002 | 0.018 |
| 7AL | right | DAN | | -0.384 | 78.703 | -3.850 | < 0.001 | 0.003 |
| 7Am | left | DAN | | -0.513 | 77.260 | -4.109 | < 0.001 | 0.002 |
| 7PC | left | DAN | | -0.387 | 77.159 | -3.837 | < 0.001 | 0.004 |

| | | | | | | | |
|---|---|---|---|---|---|---|---|
| 7PL | left | DAN | -0.423 | 77.239 | -3.455 | 0.001 | 0.010 |
| AIP | left | DAN | -0.386 | 82.445 | -3.630 | < 0.001 | 0.006 |
| AIP | right | DAN | -0.387 | 81.846 | -3.189 | 0.002 | 0.020 |
| PFt | right | DAN | -0.345 | 78.839 | -2.992 | 0.004 | 0.032 |
| 6r | right | VAN | -0.336 | 80.030 | -2.843 | 0.006 | 0.041 |
| FOP3 | right | VAN | -0.371 | 72.567 | -2.963 | 0.004 | 0.032 |
| PeEc | right | LIN | 0.521 | 70.915 | 3.823 | < 0.001 | 0.004 |
| TGv | left | LIN | 0.478 | 74.969 | 2.970 | 0.004 | 0.032 |

**Table S2 Regions that show significant differences gradient 2 scores between CB-SC groups**

| region | hemi | yeo | stream | estimate | parameter | statistic | p (uncorr) | p (corr) |
|---|---|---|---|---|---|---|---|---|
| V3B | left | VIS | dorsal | -0.563 | 60.914 | -2.526 | 0.014 | 0.027 |
| V3B | right | VIS | dorsal | -0.597 | 59.309 | -2.527 | 0.014 | 0.027 |
| V6A | left | VIS | dorsal | 0.653 | 63.222 | 3.193 | 0.002 | 0.006 |
| V6A | right | VIS | dorsal | 0.587 | 65.686 | 2.809 | 0.007 | 0.014 |
| V1 | left | VIS | evc | -1.045 | 61.54 | -4.683 | < 0.001 | < 0.001 |
| V1 | right | VIS | evc | -1.047 | 61.619 | -4.905 | < 0.001 | < 0.001 |
| V2 | left | VIS | evc | -0.758 | 63.064 | -3.394 | 0.001 | 0.003 |
| V2 | right | VIS | evc | -0.9 | 61.556 | -4.251 | < 0.001 | < 0.001 |
| V3 | left | VIS | evc | -0.741 | 63.733 | -3.42 | 0.001 | 0.003 |
| V3 | right | VIS | evc | -0.838 | 66.378 | -3.796 | < 0.001 | 0.001 |
| V4 | left | VIS | evc | -0.798 | 66.127 | -3.799 | < 0.001 | 0.001 |
| V4 | right | VIS | evc | -0.853 | 67.231 | -4.028 | < 0.001 | 0.001 |
| LO1 | right | VIS | lateral | -0.567 | 58.346 | -2.356 | 0.022 | 0.04 |
| LO2 | left | VIS | lateral | -0.617 | 61.111 | -2.605 | 0.012 | 0.023 |
| LO2 | right | VIS | lateral | -0.832 | 59.752 | -3.901 | < 0.001 | 0.001 |
| V3CD | right | VIS | lateral | -0.79 | 67.112 | -3.829 | < 0.001 | 0.001 |
| FFC | left | VIS | ventral | -0.951 | 62.885 | -4.405 | < 0.001 | < 0.001 |
| FFC | right | VIS | ventral | -1.138 | 65.146 | -5.199 | < 0.001 | < 0.001 |
| PIT | left | VIS | ventral | -0.814 | 62.607 | -3.66 | 0.001 | 0.002 |
| PIT | right | VIS | ventral | -0.934 | 62.209 | -4.126 | < 0.001 | 0.001 |
| V8 | left | VIS | ventral | -0.803 | 65.507 | -3.495 | 0.001 | 0.003 |
| V8 | right | VIS | ventral | -0.825 | 62.302 | -3.556 | 0.001 | 0.002 |
| VMV1 | left | VIS | ventral | -0.784 | 60.69 | -3.351 | 0.001 | 0.004 |
| VMV1 | right | VIS | ventral | -0.765 | 58.344 | -3.266 | 0.002 | 0.005 |
| VMV2 | left | VIS | ventral | -0.891 | 59.646 | -3.846 | < 0.001 | 0.001 |
| VMV2 | right | VIS | ventral | -0.833 | 60.159 | -3.45 | 0.001 | 0.003 |
| VMV3 | left | VIS | ventral | -0.641 | 62.681 | -2.616 | 0.011 | 0.022 |
| VMV3 | right | VIS | ventral | -0.885 | 68.304 | -3.787 | < 0.001 | 0.001 |
| VVC | left | VIS | ventral | -1.001 | 68.263 | -4.584 | < 0.001 | < 0.001 |
| VVC | right | VIS | ventral | -0.970 | 68.826 | -4.36 | < 0.001 | < 0.001 |
| PHA1 | left | VIS | | -0.946 | 57.374 | -4.771 | < 0.001 | < 0.001 |
| PHA2 | left | VIS | | -0.770 | 64.362 | -4.464 | < 0.001 | < 0.001 |
| PHA3 | left | VIS | | -0.763 | 66.413 | -4.107 | < 0.001 | 0.001 |
| PHA3 | right | VIS | | -0.708 | 57.817 | -4.028 | < 0.001 | 0.001 |
| PreS | left | VIS | | -0.761 | 57.66 | -4.267 | < 0.001 | < 0.001 |
| ProS | left | VIS | | -0.681 | 57.717 | -3.402 | 0.001 | 0.003 |
| ProS | right | VIS | | -0.56 | 53.269 | -2.906 | 0.005 | 0.011 |
| 1 | left | SMN | | 1.168 | 77.159 | 7.996 | 0 | 0 |
| 1 | right | SMN | | 1.115 | 75.171 | 7.472 | 0 | 0 |
| 2 | left | SMN | | 0.9 | 69.802 | 6.794 | 0 | 0 |
| 2 | right | SMN | | 0.909 | 68.441 | 6.311 | 0 | 0 |
| 24dd | left | SMN | | 0.738 | 61.639 | 5.251 | 0 | 0 |
| 24dd | right | SMN | | 0.744 | 65.744 | 5.575 | 0 | 0 |

| | | | | | | | |
|---|---|---|---|---|---|---|---|
| 24dv | left | SMN | 0.674 | 61.161 | 4.854 | 0 | 0 |
| 24dv | right | SMN | 0.672 | 61.832 | 5.222 | 0 | 0 |
| 3a | left | SMN | 1.014 | 74.412 | 7.076 | 0 | 0 |
| 3a | right | SMN | 1.016 | 74.216 | 7.191 | 0 | 0 |
| 3b | left | SMN | 1.013 | 73.802 | 7.305 | 0 | 0 |
| 3b | right | SMN | 1.095 | 76.101 | 7.974 | 0 | 0 |
| 4 | left | SMN | 1.056 | 78.057 | 7.596 | 0 | 0 |
| 4 | right | SMN | 1.038 | 76.113 | 7.622 | 0 | 0 |
| 43 | left | SMN | 0.764 | 68.147 | 5.277 | 0 | 0 |
| 52 | left | SMN | 0.447 | 63.068 | 2.8 | 0.007 | 0.014 |
| 52 | right | SMN | 0.589 | 58.146 | 3.895 | 0 | 0.001 |
| 5L | left | SMN | 0.911 | 72.204 | 6.283 | 0 | 0 |
| 5L | right | SMN | 0.87 | 77.589 | 5.926 | 0 | 0 |
| 5m | left | SMN | 0.894 | 69.895 | 5.806 | 0 | 0 |
| 5m | right | SMN | 0.787 | 71.41 | 4.993 | 0 | 0 |
| 6d | left | SMN | 0.929 | 73.468 | 6.529 | 0 | 0 |
| 6d | right | SMN | 0.924 | 73.838 | 6.28 | 0 | 0 |
| 6mp | left | SMN | 0.756 | 74.981 | 6.264 | 0 | 0 |
| 6mp | right | SMN | 0.812 | 73.726 | 6 | 0 | 0 |
| 6v | left | SMN | 0.72 | 61.05 | 5.139 | 0 | 0 |
| 6v | right | SMN | 0.673 | 63.79 | 4.258 | 0 | 0 |
| A1 | left | SMN | 0.578 | 62.484 | 3.589 | 0.001 | 0.002 |
| A1 | right | SMN | 0.488 | 57.749 | 2.595 | 0.012 | 0.023 |
| A4 | left | SMN | 0.596 | 63.439 | 3.836 | 0 | 0.001 |
| A4 | right | SMN | 0.586 | 63.436 | 3.693 | 0 | 0.002 |
| A5 | left | SMN | 0.594 | 75.143 | 3.893 | 0 | 0.001 |
| A5 | right | SMN | 0.651 | 75.465 | 4.149 | 0 | 0 |
| FOP2 | left | SMN | 0.661 | 59.786 | 4.243 | 0 | 0 |
| Ig | left | SMN | 0.789 | 64.489 | 4.996 | 0 | 0 |
| Ig | right | SMN | 0.684 | 60.508 | 4.076 | 0 | 0.001 |
| LBelt | left | SMN | 0.689 | 62.387 | 4.318 | 0 | 0 |
| LBelt | right | SMN | 0.638 | 61.824 | 3.832 | 0 | 0.001 |
| MBelt | left | SMN | 0.503 | 69.357 | 3.296 | 0.002 | 0.004 |
| MBelt | right | SMN | 0.481 | 63.364 | 2.977 | 0.004 | 0.009 |
| OP1 | left | SMN | 0.817 | 63.416 | 5.503 | 0 | 0 |
| OP1 | right | SMN | 0.835 | 68.158 | 5.514 | 0 | 0 |
| OP2-3 | left | SMN | 0.848 | 68.625 | 5.165 | 0 | 0 |
| OP2-3 | right | SMN | 0.622 | 58.698 | 3.567 | 0.001 | 0.002 |
| OP4 | left | SMN | 0.818 | 68.702 | 5.676 | 0 | 0 |
| OP4 | right | SMN | 0.862 | 64.453 | 5.438 | 0 | 0 |
| PBelt | left | SMN | 0.561 | 62.777 | 3.622 | 0.001 | 0.002 |
| PBelt | right | SMN | 0.559 | 64.622 | 3.52 | 0.001 | 0.002 |
| PFcm | left | SMN | 0.516 | 64.883 | 3.493 | 0.001 | 0.003 |
| RI | left | SMN | 0.789 | 66.34 | 4.956 | 0 | 0 |
| RI | right | SMN | 0.589 | 59.044 | 3.535 | 0.001 | 0.002 |
| TA2 | right | SMN | 0.497 | 67.389 | 3.298 | 0.002 | 0.004 |

| | | | | | | | | |
|---|---|---|---|---|---|---|---|---|
| IPS1 | left | DAN | dorsal | -0.708 | 59.708 | -3.515 | 0.001 | 0.003 |
| IPS1 | right | DAN | dorsal | -0.635 | 61.996 | -2.954 | 0.004 | 0.01 |
| PH | left | DAN | lateral | -0.657 | 60.513 | -3.098 | 0.003 | 0.007 |
| PH | right | DAN | lateral | -0.772 | 52.987 | -3.266 | 0.002 | 0.005 |
| 7AL | left | DAN | | 0.783 | 82.172 | 5.088 | 0 | 0 |
| 7AL | right | DAN | | 0.701 | 80.32 | 4.792 | 0 | 0 |
| 7Am | left | DAN | | 0.485 | 75.326 | 3.496 | 0.001 | 0.002 |
| 7PC | left | DAN | | 0.677 | 76.139 | 4.919 | 0 | 0 |
| 7PC | right | DAN | | 0.5 | 65.078 | 3.204 | 0.002 | 0.005 |
| IFJp | left | DAN | | -0.843 | 57.741 | -5.196 | 0 | 0 |
| IFJp | right | DAN | | -0.995 | 55.906 | -6.531 | 0 | 0 |
| IP0 | left | DAN | | -0.88 | 65.684 | -4.389 | 0 | 0 |
| IP0 | right | DAN | | -1.144 | 55.336 | -5.37 | 0 | 0 |
| LIPd | left | DAN | | -0.543 | 67.132 | -3.288 | 0.002 | 0.004 |
| LIPd | right | DAN | | -0.598 | 67.068 | -4.061 | < 0.001 | 0.001 |
| MIP | left | DAN | | -0.557 | 69.137 | -3.246 | 0.002 | 0.005 |
| MIP | right | DAN | | -0.544 | 62.568 | -3.02 | 0.004 | 0.008 |
| PEF | right | DAN | | -0.711 | 56.518 | -3.626 | 0.001 | 0.002 |
| PFt | left | DAN | | 0.52 | 64.768 | 3.691 | < 0.001 | 0.002 |
| PFt | right | DAN | | 0.443 | 55.714 | 2.962 | 0.004 | 0.01 |
| PGp | left | DAN | | -0.71 | 67.148 | -3.754 | < 0.001 | 0.001 |
| PGp | right | DAN | | -0.889 | 57.931 | -5.15 | < 0.001 | < 0.001 |
| TE2p | right | DAN | | -0.66 | 55.007 | -3.468 | 0.001 | 0.003 |
| VIP | left | DAN | | 0.635 | 81.658 | 3.558 | 0.001 | 0.002 |
| VIP | right | DAN | | 0.372 | 76.121 | 2.649 | 0.01 | 0.02 |
| 23c | left | VAN | | 0.446 | 76.398 | 4.077 | < 0.001 | 0.001 |
| 23c | right | VAN | | 0.545 | 70.956 | 4.244 | < 0.001 | < 0.001 |
| 43 | right | VAN | | 0.808 | 64.255 | 5.423 | < 0.001 | < 0.001 |
| 5mv | left | VAN | | 0.644 | 71.008 | 5.582 | < 0.001 | < 0.001 |
| 5mv | right | VAN | | 0.642 | 70.208 | 5.086 | < 0.001 | < 0.001 |
| AAIC | left | VAN | | 0.315 | 82.772 | 3.134 | 0.002 | 0.006 |
| FOP1 | left | VAN | | 0.578 | 65.926 | 4.104 | < 0.001 | 0.001 |
| FOP1 | right | VAN | | 0.396 | 60.033 | 2.456 | 0.017 | 0.032 |
| FOP3 | left | VAN | | 0.501 | 64.292 | 3.349 | 0.001 | 0.004 |
| FOP3 | right | VAN | | 0.447 | 69.002 | 2.878 | 0.005 | 0.011 |
| FOP4 | left | VAN | | 0.366 | 66.75 | 2.99 | 0.004 | 0.009 |
| FOP4 | right | VAN | | 0.313 | 73.143 | 2.365 | 0.021 | 0.039 |
| MI | left | VAN | | 0.328 | 71.844 | 2.915 | 0.005 | 0.01 |
| MI | right | VAN | | 0.333 | 71.716 | 2.964 | 0.004 | 0.009 |
| PFcm | right | VAN | | 0.51 | 61.527 | 3.32 | 0.002 | 0.004 |
| PFop | left | VAN | | 0.486 | 64.331 | 3.95 | < 0.001 | 0.001 |
| PFop | right | VAN | | 0.588 | 58.928 | 4.479 | < 0.001 | < 0.001 |
| PI | right | VAN | | 0.364 | 63.676 | 2.49 | 0.015 | 0.029 |
| PSL | left | VAN | | 0.396 | 69.974 | 3.245 | 0.002 | 0.005 |
| PoI1 | left | VAN | | 0.46 | 62.015 | 3.144 | 0.003 | 0.006 |
| PoI1 | right | VAN | | 0.484 | 59.685 | 3.425 | 0.001 | 0.003 |

| | | | | | | | |
|---|---|---|---|---|---|---|---|
| PoI2 | left | VAN | 0.534 | 59.267 | 3.877 | < 0.001 | 0.001 |
| PoI2 | right | VAN | 0.524 | 62.265 | 4.013 | < 0.001 | 0.001 |
| SCEF | left | VAN | 0.529 | 64.93 | 4.419 | < 0.001 | < 0.001 |
| SCEF | right | VAN | 0.47 | 70.971 | 3.554 | 0.001 | 0.002 |
| TPOJ1 | left | VAN | 0.458 | 73.865 | 2.759 | 0.007 | 0.015 |
| a24pr | left | VAN | 0.352 | 57.8 | 3.463 | 0.001 | 0.003 |
| a24pr | right | VAN | 0.343 | 65.527 | 2.983 | 0.004 | 0.009 |
| p24pr | left | VAN | 0.427 | 54.193 | 3.358 | 0.001 | 0.004 |
| p24pr | right | VAN | 0.505 | 57.851 | 4.512 | < 0.001 | < 0.001 |
| p32pr | left | VAN | 0.301 | 67.146 | 2.974 | 0.004 | 0.009 |
| p32pr | right | VAN | 0.32 | 63.51 | 2.454 | 0.017 | 0.032 |
| Pir | right | LIN | 0.361 | 78.951 | 2.768 | 0.007 | 0.015 |
| TF | left | LIN | -0.339 | 56.076 | -2.297 | 0.025 | 0.046 |
| TF | right | LIN | -0.336 | 64.464 | -2.67 | 0.01 | 0.019 |
| TGd | right | LIN | 0.263 | 74.822 | 2.339 | 0.022 | 0.04 |
| 11l | left | FPN | -0.279 | 64.679 | -2.321 | 0.023 | 0.043 |
| 44 | left | FPN | -0.391 | 52.578 | -3.171 | 0.003 | 0.006 |
| 8BM | left | FPN | -0.309 | 60.417 | -2.803 | 0.007 | 0.014 |
| 8BM | right | FPN | -0.279 | 55.026 | -2.296 | 0.026 | 0.046 |
| 8C | left | FPN | -0.406 | 47.161 | -3.126 | 0.003 | 0.007 |
| 8C | right | FPN | -0.441 | 49.938 | -3.482 | 0.001 | 0.003 |
| IFJa | left | FPN | -0.671 | 61.04 | -4.066 | < 0.001 | 0.001 |
| IFJa | right | FPN | -0.817 | 49.949 | -5.45 | < 0.001 | < 0.001 |
| IFSa | left | FPN | -0.656 | 59.174 | -4.882 | < 0.001 | < 0.001 |
| IFSa | right | FPN | -0.73 | 56.52 | -4.959 | < 0.001 | < 0.001 |
| IFSp | left | FPN | -0.713 | 59.882 | -4.808 | < 0.001 | < 0.001 |
| IFSp | right | FPN | -0.829 | 47.012 | -4.686 | < 0.001 | < 0.001 |
| IP1 | left | FPN | -0.822 | 52.574 | -6.027 | < 0.001 | < 0.001 |
| IP1 | right | FPN | -1.036 | 52.404 | -6.137 | < 0.001 | < 0.001 |
| IP2 | left | FPN | -0.523 | 69.085 | -4.373 | < 0.001 | < 0.001 |
| IP2 | right | FPN | -0.543 | 64.515 | -4.091 | < 0.001 | 0.001 |
| PFm | left | FPN | -0.418 | 59.825 | -4.577 | < 0.001 | < 0.001 |
| POS2 | left | FPN | -0.435 | 66.16 | -3.498 | 0.001 | 0.003 |
| a47r | left | FPN | -0.318 | 52.386 | -2.624 | 0.011 | 0.022 |
| a9-46v | left | FPN | -0.391 | 61.286 | -4.3 | < 0.001 | < 0.001 |
| i6-8 | left | FPN | -0.369 | 56.683 | -2.987 | 0.004 | 0.009 |
| i6-8 | right | FPN | -0.358 | 61.855 | -2.957 | 0.004 | 0.01 |
| p47r | left | FPN | -0.439 | 70.404 | -3.927 | < 0.001 | 0.001 |
| p47r | right | FPN | -0.489 | 63.531 | -4.279 | < 0.001 | < 0.001 |
| p9-46v | left | FPN | -0.608 | 53.404 | -4.804 | < 0.001 | < 0.001 |
| p9-46v | right | FPN | -0.342 | 53.791 | -2.653 | 0.01 | 0.021 |
| s6-8 | left | FPN | -0.228 | 65.552 | -2.346 | 0.022 | 0.04 |
| 44 | right | DMN | -0.376 | 58.596 | -3.088 | 0.003 | 0.007 |
| PFm | right | DMN | -0.32 | 58.45 | -2.989 | 0.004 | 0.009 |
| PGs | left | DMN | -0.24 | 61.51 | -2.754 | 0.008 | 0.016 |
| PGs | right | DMN | -0.279 | 59.63 | -2.934 | 0.005 | 0.01 |

| | | | | | | | |
|---|---|---|---|---|---|---|---|
| PHA1 | right | DMN | -0.671 | 56.312 | -3.719 | < 0.001 | 0.002 |
| PHA2 | right | DMN | -0.526 | 57.245 | -3.187 | 0.002 | 0.006 |
| POS1 | left | DMN | -0.444 | 67.579 | -3.319 | 0.001 | 0.004 |
| POS1 | right | DMN | -0.389 | 66.295 | -3.019 | 0.004 | 0.008 |
| POS2 | right | DMN | -0.341 | 61.515 | -2.732 | 0.008 | 0.017 |
| PreS | right | DMN | -0.613 | 52.725 | -3.953 | < 0.001 | 0.001 |
| RSC | left | DMN | -0.336 | 69.896 | -2.786 | 0.007 | 0.014 |
| STGa | right | DMN | 0.499 | 66.973 | 4.12 | < 0.001 | 0.001 |
| STSda | right | DMN | 0.424 | 72.594 | 3.082 | 0.003 | 0.007 |
| STSdp | left | DMN | 0.549 | 71.6 | 3.606 | 0.001 | 0.002 |
| STSdp | right | DMN | 0.431 | 71.202 | 3.091 | 0.003 | 0.007 |
| STSva | right | DMN | 0.331 | 68.213 | 3.237 | 0.002 | 0.005 |
| STSvp | left | DMN | 0.448 | 60.457 | 3.546 | 0.001 | 0.002 |
| STSvp | right | DMN | 0.5 | 64.673 | 4.16 | < 0.001 | < 0.001 |
| TE1a | right | DMN | 0.22 | 65.458 | 2.373 | 0.021 | 0.039 |
| TE1m | right | DMN | 0.254 | 71.436 | 2.6 | 0.011 | 0.022 |
| a47r | right | DMN | -0.389 | 52.509 | -3.111 | 0.003 | 0.007 |
| s6-8 | right | DMN | -0.241 | 64.881 | -2.419 | 0.018 | 0.035 |

**Table S3 Regions that show significant differences gradient 3 scores between CB-SC groups**

| region | hemi | yeo | stream | estimate | parameter | statistic | p (uncorr) | p (corr) |
|---|---|---|---|---|---|---|---|---|
| V3A | left | VIS | dorsal | -0.802 | 76.887 | -6.8 | < 0.001 | <0.001 |
| V3A | right | VIS | dorsal | -0.671 | 76.095 | -5.456 | < 0.001 | <0.001 |
| V3B | left | VIS | dorsal | -0.717 | 78.274 | -4.086 | < 0.001 | 0.002 |
| V3B | right | VIS | dorsal | -0.707 | 82.891 | -3.947 | < 0.001 | 0.002 |
| V6 | left | VIS | dorsal | -0.461 | 82.382 | -3.286 | 0.001 | 0.015 |
| V6 | right | VIS | dorsal | -0.382 | 82.917 | -2.942 | 0.004 | 0.037 |
| V6A | left | VIS | dorsal | -0.901 | 82.883 | -5.918 | < 0.001 | <0.001 |
| V6A | right | VIS | dorsal | -0.887 | 82.799 | -6.388 | < 0.001 | <0.001 |
| V7 | left | VIS | dorsal | -0.849 | 82.036 | -4.451 | < 0.001 | 0.001 |
| V7 | right | VIS | dorsal | -0.75 | 82.994 | -5.019 | < 0.001 | <0.001 |
| V2 | left | VIS | evc | -0.417 | 70.749 | -3.538 | 0.001 | 0.009 |
| V3 | left | VIS | evc | -0.677 | 74.911 | -5.386 | < 0.001 | <0.001 |
| V3 | right | VIS | evc | -0.615 | 80.007 | -5.531 | < 0.001 | <0.001 |
| V4 | left | VIS | evc | -0.805 | 76.29 | -6.222 | < 0.001 | <0.001 |
| V4 | right | VIS | evc | -0.809 | 77.34 | -6.173 | < 0.001 | <0.001 |
| LO1 | left | VIS | lateral | -0.69 | 71.127 | -4.164 | < 0.001 | 0.001 |
| LO2 | left | VIS | lateral | -0.578 | 77.054 | -5.054 | < 0.001 | <0.001 |
| LO2 | right | VIS | lateral | -0.563 | 82.16 | -4.559 | < 0.001 | <0.001 |
| V3CD | left | VIS | lateral | -0.591 | 82.224 | -3.834 | < 0.001 | 0.003 |
| V4t | left | VIS | lateral | -0.415 | 82.54 | -3.402 | 0.001 | 0.011 |
| V4t | right | VIS | lateral | -0.407 | 82.888 | -3.241 | 0.002 | 0.017 |
| PIT | left | VIS | ventral | -0.686 | 75.162 | -4.77 | < 0.001 | <0.001 |
| PIT | right | VIS | ventral | -0.485 | 82.446 | -3.727 | < 0.001 | 0.005 |
| V8 | left | VIS | ventral | -0.654 | 81.328 | -4.862 | < 0.001 | <0.001 |
| V8 | right | VIS | ventral | -0.582 | 80.01 | -4.147 | < 0.001 | 0.001 |
| VMV3 | right | VIS | ventral | -0.476 | 76.647 | -3.204 | 0.002 | 0.019 |
| DVT | left | VIS | | -0.562 | 81.685 | -3.024 | 0.003 | 0.03 |
| PHA2 | left | VIS | | 0.509 | 82.525 | 3.469 | 0.001 | 0.009 |
| IFJp | right | DAN | | 0.658 | 82.945 | 3.369 | 0.001 | 0.012 |
| IP0 | right | DAN | | 0.665 | 81.71 | 3.13 | 0.002 | 0.023 |
| PGp | right | DAN | | 0.766 | 82.741 | 4.5 | < 0.001 | <0.001 |
| TE2p | right | DAN | | 0.901 | 82.618 | 4.538 | < 0.001 | <0.001 |
| H | left | LIN | | 0.438 | 81.151 | 3.652 | < 0.001 | 0.006 |
| H | right | LIN | | 0.501 | 81.624 | 3.78 | < 0.001 | 0.004 |
| PeEc | right | LIN | | 0.712 | 76.749 | 4.623 | < 0.001 | <0.001 |
| TF | left | LIN | | 0.71 | 77.996 | 4.682 | < 0.001 | <0.001 |
| TF | right | LIN | | 0.655 | 80.531 | 3.968 | < 0.001 | 0.002 |
| IFJa | right | FPN | | 0.801 | 82.772 | 3.922 | < 0.001 | 0.003 |
| IFSa | right | FPN | | 0.58 | 82.478 | 3.028 | 0.003 | 0.03 |
| IP1 | right | FPN | | 0.855 | 80.633 | 4.479 | < 0.001 | <0.001 |
| p24 | right | DMN | | -0.533 | 79.176 | -3.477 | 0.001 | 0.009 |

**Table S4 Summary of the regions that show significant differences in gradient scores between CB and SC groups across 3 gradients.** For G1, G2, G3: If 1, then region shows a significant group difference.

| region | hemi | stream | G1 | G2 | G3 |
|---|---|---|---|---|---|
| V3A | left | dorsal | 1 | 0 | 1 |
| V3A | right | dorsal | 1 | 0 | 1 |
| V3B | left | dorsal | 1 | 1 | 1 |
| V3B | right | dorsal | 1 | 1 | 1 |
| V7 | left | dorsal | 1 | 0 | 1 |
| V7 | right | dorsal | 0 | 0 | 1 |
| V6A | left | dorsal | 0 | 1 | 1 |
| V6A | right | dorsal | 0 | 1 | 1 |
| V6 | left | dorsal | 0 | 0 | 1 |
| V6 | right | dorsal | 0 | 0 | 1 |
| V1 | left | evc | 1 | 1 | 0 |
| V1 | right | evc | 1 | 1 | 0 |
| V2 | left | evc | 1 | 1 | 1 |
| V2 | right | evc | 1 | 1 | 0 |
| V3 | left | evc | 1 | 1 | 1 |
| V3 | right | evc | 1 | 1 | 1 |
| V4 | left | evc | 1 | 1 | 1 |
| V4 | right | evc | 1 | 1 | 1 |
| LO1 | left | lateral | 1 | 0 | 1 |
| LO1 | right | lateral | 1 | 1 | 0 |
| LO2 | left | lateral | 1 | 1 | 1 |
| LO2 | right | lateral | 1 | 1 | 1 |
| V3CD | left | lateral | 1 | 0 | 1 |
| V3CD | right | lateral | 1 | 1 | 0 |
| V4t | left | lateral | 1 | 0 | 1 |
| V4t | right | lateral | 1 | 0 | 1 |
| FFC | left | ventral | 1 | 1 | 0 |
| FFC | right | ventral | 1 | 1 | 0 |
| PIT | left | ventral | 1 | 1 | 1 |
| PIT | right | ventral | 1 | 1 | 1 |
| V8 | left | ventral | 1 | 1 | 1 |
| V8 | right | ventral | 1 | 1 | 1 |
| VMV1 | left | ventral | 1 | 1 | 0 |
| VMV1 | right | ventral | 1 | 1 | 0 |
| VMV2 | left | ventral | 0 | 1 | 0 |
| VMV2 | right | ventral | 1 | 1 | 0 |
| VMV3 | left | ventral | 1 | 1 | 0 |
| VMV3 | right | ventral | 1 | 1 | 1 |
| VVC | left | ventral | 1 | 1 | 0 |
| VVC | right | ventral | 1 | 1 | 0 |
| PHA1 | left | ventral | 0 | 1 | 0 |

| | | | | | |
|---|---|---|---|---|---|
| PHA2 | left | ventral | 0 | 1 | 1 |
| PHA3 | left | ventral | 0 | 1 | 0 |
| PHA3 | right | ventral | 0 | 1 | 0 |
| PreS | left | ventral | 0 | 1 | 0 |
| ProS | left | ventral | 0 | 1 | 0 |
| ProS | right | ventral | 0 | 1 | 0 |
| DVT | left | ventral | 0 | 0 | 1 |

**Table S5 | Group differences in community-based profiles of the first three gradients.**

| yeo | gradient | estimate | parameter | statistic | p (uncorr) | CI (low) | CI (high) | p (corr) |
|---|---|---|---|---|---|---|---|---|
| VN | 1 | -0.392 | 51.925 | -4.943 | 0 | -0.551 | -0.233 | 0 |
| SMN | 1 | 0.219 | 51.499 | 3.897 | 0 | 0.106 | 0.331 | 0.001 |
| DAN | 1 | 0.092 | 46.529 | 0.82 | 0.417 | -0.134 | 0.317 | 0.486 |
| VAN | 1 | 0.2 | 51.116 | 2.693 | 0.01 | 0.051 | 0.35 | 0.022 |
| LN | 1 | -0.175 | 27.903 | -1.032 | 0.311 | -0.523 | 0.172 | 0.435 |
| FPN | 1 | 0.104 | 53.84 | 1.128 | 0.264 | -0.08 | 0.288 | 0.435 |
| DMN | 1 | -0.044 | 97.997 | -0.429 | 0.669 | -0.249 | 0.161 | 0.669 |
| VN | 2 | 0.575 | 49.015 | 5.446 | 0 | 0.363 | 0.787 | 0 |
| SMN | 2 | -0.7 | 42.367 | -12.791 | 0 | -0.811 | -0.59 | 0 |
| DAN | 2 | 0.137 | 34.281 | 0.878 | 0.386 | -0.18 | 0.453 | 0.386 |
| VAN | 2 | -0.332 | 36.324 | -4.889 | 0 | -0.47 | -0.194 | 0 |
| LN | 2 | 0.062 | 19.275 | 0.888 | 0.386 | -0.083 | 0.206 | 0.386 |
| FPN | 2 | 0.318 | 39.347 | 4.759 | 0 | 0.183 | 0.453 | 0 |
| DMN | 2 | 0.058 | 69.038 | 0.972 | 0.334 | -0.061 | 0.176 | 0.386 |
| VN | 3 | 0.304 | 52.754 | 3.999 | 0 | 0.152 | 0.457 | 0.001 |
| SMN | 3 | -0.055 | 54.997 | -0.971 | 0.336 | -0.167 | 0.058 | 0.784 |
| DAN | 3 | -0.109 | 47.652 | -0.682 | 0.498 | -0.431 | 0.213 | 0.787 |
| VAN | 3 | -0.012 | 50.436 | -0.112 | 0.911 | -0.235 | 0.21 | 0.912 |
| LN | 3 | -0.255 | 27.762 | -2.153 | 0.04 | -0.498 | -0.012 | 0.141 |
| FPN | 3 | -0.074 | 53.088 | -0.584 | 0.562 | -0.328 | 0.18 | 0.787 |
| DMN | 3 | -0.011 | 98 | -0.111 | 0.912 | -0.212 | 0.19 | 0.912 |

**Table S6 | Estimated marginal means of VN distance to the other 6 yeo networks in G1 per group and network.** CB = congenitally blind, SC = sighted control.

| group | yeo | emmean | SE | df | lower.CL | upper.CL |
|---|---|---|---|---|---|---|
| SC | SMN | 0.319 | 0.083 | 281.246 | 0.155 | 0.483 |
| CB | SMN | 0.975 | 0.086 | 282.011 | 0.805 | 1.145 |
| SC | DAN | 0.333 | 0.083 | 281.246 | 0.169 | 0.497 |
| CB | DAN | 0.344 | 0.086 | 282.011 | 0.174 | 0.514 |
| SC | VAN | 0.294 | 0.083 | 281.246 | 0.129 | 0.458 |
| CB | VAN | 0.702 | 0.086 | 282.011 | 0.532 | 0.872 |
| SC | LN | 0.963 | 0.083 | 281.246 | 0.798 | 1.127 |
| CB | LN | 0.764 | 0.086 | 282.011 | 0.594 | 0.934 |
| SC | FPN | 1.003 | 0.083 | 281.246 | 0.839 | 1.167 |
| CB | FPN | 0.54 | 0.086 | 282.011 | 0.37 | 0.710 |
| SC | DMN | 1.295 | 0.083 | 281.246 | 1.13 | 1.459 |
| CB | DMN | 0.944 | 0.086 | 282.011 | 0.774 | 1.114 |

**Table S7 | Results of post hoc tests assessing the effect of group on distance of VN to the other 6 yeo networks for each network in G1.**

| yeo | estimate | SE | df | t-ratio | p-value |
|---|---|---|---|---|---|
| **SMN** | **-0.656** | **0.12** | **281.43** | **-5.464** | **< 0.001** |
| DAN | -0.011 | 0.12 | 281.43 | -0.094 | 0.925 |
| **VAN** | **-0.408** | **0.12** | **281.43** | **-3.399** | **0.001** |
| LN | 0.199 | 0.12 | 281.43 | 1.657 | 0.099 |
| **FPN** | **0.463** | **0.12** | **281.43** | **3.858** | **< 0.001** |
| **DMN** | **0.351** | **0.12** | **281.43** | **2.92** | **0.004** |

**Table S8 | Estimated marginal means of VN distance to the other 6 yeo networks in G2 per group and network.** CB = congenitally blind, SC = sighted control.

| group | yeo | emmean | SE | df | lower.CL | upper.CL |
|---|---|---|---|---|---|---|
| SC | SMN | 1.409 | 0.142 | 100.574 | 1.127 | 1.692 |
| CB | SMN | 2.679 | 0.147 | 100.657 | 2.386 | 2.971 |
| SC | DAN | 0.783 | 0.142 | 100.574 | 0.501 | 1.066 |
| CB | DAN | 1.215 | 0.147 | 100.657 | 0.923 | 1.508 |
| SC | VAN | 1.384 | 0.142 | 100.574 | 1.101 | 1.667 |
| CB | VAN | 2.285 | 0.147 | 100.657 | 1.993 | 2.577 |
| SC | LN | 1 | 0.142 | 100.574 | 0.717 | 1.283 |
| CB | LN | 1.506 | 0.147 | 100.657 | 1.214 | 1.798 |
| SC | FPN | 1.004 | 0.142 | 100.574 | 0.722 | 1.287 |
| CB | FPN | 1.255 | 0.147 | 100.657 | 0.963 | 1.548 |
| SC | DMN | 0.937 | 0.142 | 100.574 | 0.655 | 1.220 |
| CB | DMN | 1.449 | 0.147 | 100.657 | 1.156 | 1.741 |

**Table 9 | Results of post hoc tests assessing the effect of group on distance of VN to the other 6 yeo networks for each network in G2.**

| yeo | estimate | SE | df | t-ratio | p-value |
|---|---|---|---|---|---|
| **SMN** | **-1.269** | **0.205** | **100.594** | **-6.190** | **< 0.001** |
| **DAN** | **-0.432** | **0.205** | **100.594** | **-2.108** | **0.038** |
| **VAN** | **-0.901** | **0.205** | **100.594** | **-4.394** | **< 0.001** |
| **LN** | **-0.506** | **0.205** | **100.594** | **-2.468** | **0.015** |
| FPN | -0.251 | 0.205 | 100.594 | -1.225 | 0.224 |
| **DMN** | **-0.511** | **0.205** | **100.594** | **-2.494** | **0.014** |

**Table S10 | Estimated marginal means of VN distance to the other 6 yeo networks in G3 per group and network.** CB = congenitally blind, SC = sighted control.

| group | yeo | emmean | SE | df | lower.CL | upper.CL |
|---|---|---|---|---|---|---|
| SC | SMN | 0.333 | 0.057 | 341.77 | 0.220 | 0.446 |
| CB | SMN | 0.263 | 0.059 | 342.611 | 0.146 | 0.379 |
| SC | DAN | 1.185 | 0.057 | 341.77 | 1.072 | 1.297 |
| CB | DAN | 0.773 | 0.059 | 342.611 | 0.656 | 0.890 |
| SC | VAN | 0.873 | 0.057 | 341.77 | 0.761 | 0.986 |
| CB | VAN | 0.609 | 0.059 | 342.611 | 0.493 | 0.726 |
| SC | LN | 0.263 | 0.057 | 341.77 | 0.150 | 0.376 |
| CB | LN | 0.454 | 0.059 | 342.611 | 0.338 | 0.571 |
| SC | FPN | 1.442 | 0.057 | 341.77 | 1.329 | 1.555 |
| CB | FPN | 1.065 | 0.059 | 342.611 | 0.948 | 1.181 |
| SC | DMN | 0.28 | 0.057 | 341.77 | 0.167 | 0.393 |
| CB | DMN | 0.271 | 0.059 | 342.611 | 0.154 | 0.387 |

**Table S11| Results of post hoc tests assessing the effect of group on distance of VN to the other 6 yeo networks for each network in G3.**

| yeo | estimate | SE | df | t-ratio | p-value |
|---|---|---|---|---|---|
| SMN | 0.07 | 0.082 | 341.972 | 0.852 | 0.395 |
| **DAN** | **0.412** | **0.082** | **341.972** | **4.993** | **< 0.001** |
| **VAN** | **0.264** | **0.082** | **341.972** | **3.202** | **0.001** |
| **LN** | **-0.191** | **0.082** | **341.972** | **-2.32** | **0.021** |
| **FPN** | **0.377** | **0.082** | **341.972** | **4.575** | **< 0.001** |
| DMN | 0.009 | 0.082 | 341.972 | 0.113 | 0.910 |

### Table S12| Occurence of the rank orders across 10000 bootstraps and 3 gradients

| G1 | | | | G2 | | | | G3 | | | |
|---|---|---|---|---|---|---|---|---|---|---|---|
| SC | | CB | | SC | | CB | | SC | | CB | |
| rank combination | frequency | rank combination | frequency | rank combination | frequency | rank combination | frequency | rank combination | frequency | rank combination | frequency |
| V1-V2-V3-V4 | 4265 | V1-V2-V3-V4 | 514 | V1-V2-V3-V4 | 6147 | V1-V2-V3-V4 | 5540 | V1-V2-V3-V4 | 52 | V1-V2-V3-V4 | 1506 |
| V1-V2-V4-V3 | 2840 | V1-V2-V4-V3 | 1213 | V1-V2-V4-V3 | 3400 | V1-V2-V4-V3 | 409 | V1-V2-V4-V3 | 7 | V1-V2-V4-V3 | 104 |
| V1-V3-V2-V4 | 58 | V1-V3-V2-V4 | 0 | V1-V3-V2-V4 | 85 | V1-V3-V2-V4 | 198 | V1-V3-V2-V4 | 31 | V1-V3-V2-V4 | 16 |
| V1-V3-V4-V2 | 197 | V1-V3-V4-V2 | 86 | V1-V3-V4-V2 | 228 | V1-V3-V4-V2 | 106 | V1-V3-V4-V2 | 2 | V1-V3-V4-V2 | 0 |
| V1-V4-V2-V3 | 7 | V1-V4-V2-V3 | 0 | V1-V4-V2-V3 | 16 | V1-V4-V2-V3 | 184 | V1-V4-V2-V3 | 5 | V1-V4-V2-V3 | 0 |
| V1-V4-V3-V2 | 164 | V1-V4-V3-V2 | 2 | V1-V4-V3-V2 | 87 | V1-V4-V3-V2 | 229 | V1-V4-V3-V2 | 4 | V1-V4-V3-V2 | 0 |
| V2-V1-V3-V4 | 1954 | V2-V1-V3-V4 | 863 | V2-V1-V3-V4 | 35 | V2-V1-V3-V4 | 1654 | V2-V1-V3-V4 | 161 | V2-V1-V3-V4 | 6942 |
| V2-V1-V4-V3 | 234 | V2-V1-V4-V3 | 2913 | V2-V1-V4-V3 | 1 | V2-V1-V4-V3 | 387 | V2-V1-V4-V3 | 15 | V2-V1-V4-V3 | 613 |
| V2-V3-V1-V4 | 7 | V2-V3-V1-V4 | 0 | V2-V3-V1-V4 | 0 | V2-V3-V1-V4 | 33 | V2-V3-V1-V4 | 104 | V2-V3-V1-V4 | 2 |
| V2-V3-V4-V1 | 4 | V2-V3-V4-V1 | 192 | V2-V3-V4-V1 | 0 | V2-V3-V4-V1 | 16 | V2-V3-V4-V1 | 7 | V2-V3-V4-V1 | 0 |
| V2-V4-V1-V3 | 1 | V2-V4-V1-V3 | 0 | V2-V4-V1-V3 | 0 | V2-V4-V1-V3 | 57 | V2-V4-V1-V3 | 49 | V2-V4-V1-V3 | 0 |
| V2-V4-V3-V1 | 1 | V2-V4-V3-V1 | 4 | V2-V4-V3-V1 | 0 | V2-V4-V3-V1 | 22 | V2-V4-V3-V1 | 4 | V2-V4-V3-V1 | 0 |
| V3-V1-V2-V4 | 215 | V3-V1-V2-V4 | 117 | V3-V1-V2-V4 | 0 | V3-V1-V2-V4 | 297 | V3-V1-V2-V4 | 1242 | V3-V1-V2-V4 | 568 |
| V3-V1-V4-V2 | 8 | V3-V1-V4-V2 | 2540 | V3-V1-V4-V2 | 0 | V3-V1-V4-V2 | 111 | V3-V1-V4-V2 | 14 | V3-V1-V4-V2 | 2 |
| V3-V2-V1-V4 | 14 | V3-V2-V1-V4 | 0 | V3-V2-V1-V4 | 0 | V3-V2-V1-V4 | 241 | V3-V2-V1-V4 | 932 | V3-V2-V1-V4 | 195 |
| V3-V2-V4-V1 | 7 | V3-V2-V4-V1 | 654 | V3-V2-V4-V1 | 0 | V3-V2-V4-V1 | 99 | V3-V2-V4-V1 | 5 | V3-V2-V4-V1 | 0 |
| V3-V4-V1-V2 | 0 | V3-V4-V1-V2 | 0 | V3-V4-V1-V2 | 0 | V3-V4-V1-V2 | 60 | V3-V4-V1-V2 | 110 | V3-V4-V1-V2 | 0 |
| V3-V4-V2-V1 | 1 | V3-V4-V2-V1 | 3 | V3-V4-V2-V1 | 0 | V3-V4-V2-V1 | 265 | V3-V4-V2-V1 | 82 | V3-V4-V2-V1 | 0 |
| V4-V1-V2-V3 | 13 | V4-V1-V2-V3 | 88 | V4-V1-V2-V3 | 0 | V4-V1-V2-V3 | 35 | V4-V1-V2-V3 | 1195 | V4-V1-V2-V3 | 18 |
| V4-V1-V3-V2 | 4 | V4-V1-V3-V2 | 534 | V4-V1-V3-V2 | 1 | V4-V1-V3-V2 | 14 | V4-V1-V3-V2 | 129 | V4-V1-V3-V2 | 15 |
| V4-V2-V1-V3 | 2 | V4-V2-V1-V3 | 0 | V4-V2-V1-V3 | 0 | V4-V2-V1-V3 | 12 | V4-V2-V1-V3 | 1607 | V4-V2-V1-V3 | 18 |
| V4-V2-V3-V1 | 1 | V4-V2-V3-V1 | 276 | V4-V2-V3-V1 | 0 | V4-V2-V3-V1 | 23 | V4-V2-V3-V1 | 359 | V4-V2-V3-V1 | 0 |
| V4-V3-V1-V2 | 1 | V4-V3-V1-V2 | 0 | V4-V3-V1-V2 | 0 | V4-V3-V1-V2 | 5 | V4-V3-V1-V2 | 1550 | V4-V3-V1-V2 | 1 |
| V4-V3-V2-V1 | 2 | V4-V3-V2-V1 | 1 | V4-V3-V2-V1 | 0 | V4-V3-V2-V1 | 3 | V4-V3-V2-V1 | 2334 | V4-V3-V2-V1 | 0 |

### Table S13| Occurrence of the rank orders across subjects and 3 gradients

| G1 | | | | G2 | | | | G3 | | | |
|---|---|---|---|---|---|---|---|---|---|---|---|
| SC | | CB | | SC | | CB | | SC | | CB | |
| rank combination | frequency | rank combination | frequency | rank combination | frequency | rank combination | frequency | rank combination | frequency | rank combination | frequency |
| V1-V2-V3-V4 | 14 | V1-V2-V3-V4 | 7 | V1-V2-V3-V4 | 10 | V1-V2-V3-V4 | 18 | V1-V2-V3-V4 | 8 | V1-V2-V3-V4 | 13 |
| V1-V2-V4-V3 | 5 | V1-V2-V4-V3 | 1 | V1-V2-V4-V3 | 1 | V1-V2-V4-V3 | 10 | V1-V2-V4-V3 | 0 | V1-V2-V4-V3 | 0 |
| V1-V3-V2-V4 | 0 | V1-V3-V2-V4 | 0 | V1-V3-V2-V4 | 2 | V1-V3-V2-V4 | 2 | V1-V3-V2-V4 | 2 | V1-V3-V2-V4 | 0 |
| V1-V3-V4-V2 | 2 | V1-V3-V4-V2 | 2 | V1-V3-V4-V2 | 1 | V1-V3-V4-V2 | 4 | V1-V3-V4-V2 | 0 | V1-V3-V4-V2 | 0 |
| V1-V4-V2-V3 | 0 | V1-V4-V2-V3 | 0 | V1-V4-V2-V3 | 1 | V1-V4-V2-V3 | 0 | V1-V4-V2-V3 | 0 | V1-V4-V2-V3 | 1 |
| V1-V4-V3-V2 | 3 | V1-V4-V3-V2 | 2 | V1-V4-V3-V2 | 1 | V1-V4-V3-V2 | 1 | V1-V4-V3-V2 | 1 | V1-V4-V3-V2 | 0 |
| V2-V1-V3-V4 | 7 | V2-V1-V3-V4 | 7 | V2-V1-V3-V4 | 7 | V2-V1-V3-V4 | 0 | V2-V1-V3-V4 | 5 | V2-V1-V3-V4 | 16 |
| V2-V1-V4-V3 | 1 | V2-V1-V4-V3 | 6 | V2-V1-V4-V3 | 3 | V2-V1-V4-V3 | 1 | V2-V1-V4-V3 | 2 | V2-V1-V4-V3 | 1 |
| V2-V3-V1-V4 | 0 | V2-V3-V1-V4 | 0 | V2-V3-V1-V4 | 1 | V2-V3-V1-V4 | 0 | V2-V3-V1-V4 | 1 | V2-V3-V1-V4 | 1 |
| V2-V3-V4-V1 | 0 | V2-V3-V4-V1 | 3 | V2-V3-V4-V1 | 1 | V2-V3-V4-V1 | 1 | V2-V3-V4-V1 | 0 | V2-V3-V4-V1 | 0 |
| V2-V4-V1-V3 | 0 | V2-V4-V1-V3 | 0 | V2-V4-V1-V3 | 1 | V2-V4-V1-V3 | 0 | V2-V4-V1-V3 | 0 | V2-V4-V1-V3 | 0 |
| V2-V4-V3-V1 | 1 | V2-V4-V3-V1 | 0 | V2-V4-V3-V1 | 0 | V2-V4-V3-V1 | 0 | V2-V4-V3-V1 | 1 | V2-V4-V3-V1 | 1 |
| V3-V1-V2-V4 | 2 | V3-V1-V2-V4 | 0 | V3-V1-V2-V4 | 3 | V3-V1-V2-V4 | 0 | V3-V1-V2-V4 | 2 | V3-V1-V2-V4 | 0 |
| V3-V1-V4-V2 | 1 | V3-V1-V4-V2 | 1 | V3-V1-V4-V2 | 0 | V3-V1-V4-V2 | 0 | V3-V1-V4-V2 | 0 | V3-V1-V4-V2 | 0 |
| V3-V2-V1-V4 | 0 | V3-V2-V1-V4 | 0 | V3-V2-V1-V4 | 2 | V3-V2-V1-V4 | 0 | V3-V2-V1-V4 | 1 | V3-V2-V1-V4 | 2 |
| V3-V2-V4-V1 | 0 | V3-V2-V4-V1 | 2 | V3-V2-V4-V1 | 2 | V3-V2-V4-V1 | 1 | V3-V2-V4-V1 | 0 | V3-V2-V4-V1 | 0 |
| V3-V4-V1-V2 | 0 | V3-V4-V1-V2 | 0 | V3-V4-V1-V2 | 0 | V3-V4-V1-V2 | 0 | V3-V4-V1-V2 | 0 | V3-V4-V1-V2 | 0 |
| V3-V4-V2-V1 | 0 | V3-V4-V2-V1 | 1 | V3-V4-V2-V1 | 1 | V3-V4-V2-V1 | 1 | V3-V4-V2-V1 | 1 | V3-V4-V2-V1 | 0 |
| V4-V1-V2-V3 | 0 | V4-V1-V2-V3 | 0 | V4-V1-V2-V3 | 1 | V4-V1-V2-V3 | 1 | V4-V1-V2-V3 | 3 | V4-V1-V2-V3 | 0 |
| V4-V1-V3-V2 | 1 | V4-V1-V3-V2 | 1 | V4-V1-V3-V2 | 0 | V4-V1-V3-V2 | 1 | V4-V1-V3-V2 | 0 | V4-V1-V3-V2 | 2 |
| V4-V2-V1-V3 | 0 | V4-V2-V1-V3 | 2 | V4-V2-V1-V3 | 0 | V4-V2-V1-V3 | 0 | V4-V2-V1-V3 | 1 | V4-V2-V1-V3 | 0 |
| V4-V2-V3-V1 | 1 | V4-V2-V3-V1 | 3 | V4-V2-V3-V1 | 0 | V4-V2-V3-V1 | 0 | V4-V2-V3-V1 | 1 | V4-V2-V3-V1 | 0 |
| V4-V3-V1-V2 | 1 | V4-V3-V1-V2 | 0 | V4-V3-V1-V2 | 0 | V4-V3-V1-V2 | 0 | V4-V3-V1-V2 | 3 | V4-V3-V1-V2 | 1 |
| V4-V3-V2-V1 | 5 | V4-V3-V2-V1 | 3 | V4-V3-V2-V1 | 3 | V4-V3-V2-V1 | 3 | V4-V3-V2-V1 | 12 | V4-V3-V2-V1 | 3 |

**Table S14 | Estimated marginal means for minimum distances in G1 as a function of group and ROI.**

| ROI | group | response | SE | df | lower.CL | upper.CL |
|---|---|---|---|---|---|---|
| V1 | CB | 0.1102 | 0.0240 | 199 | 0.0718 | 0.1693 |
| V1 | SC | 0.3728 | 0.0783 | 199 | 0.2464 | 0.5640 |
| V2 | CB | 0.0983 | 0.0214 | 199 | 0.0640 | 0.1509 |
| V2 | SC | 0.1422 | 0.0299 | 199 | 0.0940 | 0.2151 |
| V3 | CB | 0.0636 | 0.0138 | 199 | 0.0414 | 0.0976 |
| V3 | SC | 0.1207 | 0.0254 | 199 | 0.0798 | 0.1827 |
| V4 | CB | 0.1232 | 0.0268 | 199 | 0.0802 | 0.1891 |
| V4 | SC | 0.1408 | 0.0296 | 199 | 0.0931 | 0.2131 |

**Table S15 | Results of post hoc tests assessing the effect of group on minimum distances in G1 within each ROI.**

| ROI | ratio | SE | df | t-ratio | p-value |
|---|---|---|---|---|---|
| V1 | 0.543 | 0.186 | 207, 1 | -1.778 | 0.308 |
| V2 | 0.683 | 0.235 | 207, 1 | -1.109 | 0.806 |
| V3 | 1.369 | 0.470 | 207, 1 | 0.916 | 0.806 |
| V4 | 1.447 | 0.497 | 207, 1 | 1.076 | 0.806 |

**Table S16| Estimated marginal means for minimum distances in G2 as a function of group and ROI.**

| ROI | group | response | SE | df | lower.CL | upper.CL |
|---|---|---|---|---|---|---|
| V1 | CB | 0.0702 | 0.01690 | 183 | 0.0437 | 0.1129 |
| V1 | SC | 0.1250 | 0.02904 | 183 | 0.0790 | 0.1976 |
| V2 | CB | 0.0592 | 0.01425 | 183 | 0.0368 | 0.0952 |
| V2 | SC | 0.0624 | 0.01451 | 183 | 0.0395 | 0.0987 |
| V3 | CB | 0.0867 | 0.02088 | 183 | 0.0539 | 0.1395 |
| V3 | SC | 0.0377 | 0.00875 | 183 | 0.0238 | 0.0596 |
| V4 | CB | 0.1445 | 0.03478 | 183 | 0.0899 | 0.2323 |
| V4 | SC | 0.0578 | 0.01344 | 183 | 0.0366 | 0.0915 |

**Table S17 | Results of post hoc tests assessing the effect of group on minimum distances in G2 within each ROI.**

| ROI | ratio | SE | df | t-ratio | p-value |
|---|---|---|---|---|---|
| **V1** | **0.296** | **0.089** | **199, 1** | **-4.029** | **< 0.001** |
| V2 | 0.691 | 0.209 | 199, 1 | -1.221 | 0.447 |
| V3 | 0.527 | 0.159 | 199, 1 | -2.121 | 0.106 |
| V4 | 0.875 | 0.264 | 199, 1 | -0.443 | 0.658 |

**Table S18 | Estimated marginal means for minimum distances in G3 as a function of group and ROI.**

| group | ROI | response | SE | df | lower.CL | upper.CL |
|---|---|---|---|---|---|---|
| CB | V1 | 0.070 | 0.017 | 183.136 | 0.044 | 0.113 |
| SC | V1 | 0.125 | 0.029 | 183.136 | 0.079 | 0.198 |
| CB | V2 | 0.059 | 0.014 | 183.136 | 0.037 | 0.095 |
| SC | V2 | 0.062 | 0.015 | 183.136 | 0.039 | 0.099 |
| CB | V3 | 0.087 | 0.021 | 183.136 | 0.054 | 0.139 |
| SC | V3 | 0.038 | 0.009 | 183.136 | 0.024 | 0.060 |
| CB | V4 | 0.144 | 0.035 | 183.136 | 0.090 | 0.232 |
| SC | V4 | 0.058 | 0.013 | 183.136 | 0.037 | 0.091 |

**Table S19 | Results of post hoc tests assessing the effect of group on minimum distances in G3 within each ROI.**

| ROI | ratio | SE | df | t-ratio | p-value |
|---|---|---|---|---|---|
| V1 | 0.562 | 0.188 | 183, 1 | -1.723 | 0.173 |
| V2 | 0.949 | 0.317 | 183, 1 | -0.158 | 0.875 |
| **V3** | **2.302** | **0.77** | **183, 1** | **2.492** | **0.041** |
| **V4** | **2.499** | **0.836** | **183, 1** | **2.737** | **0.027** |

**Table S20 | Anova and Tukey-Kramer post-hoc tests of the mean absolute coupling values across groups and gradients.**

| | Sum Sq. | df | Mean Sq. | F | p-value |
|---|---|---|---|---|---|
| Group | 0.03 | 1 | 0.03 | 2.81 | 0.090 |
| Gradient | 7.54 | 2 | 3.77 | 276.01 | < 0.001 |
| Group X Gradient | 3.45 | 2 | 1.72 | 63.62 | < 0.001 |
| Error | 29.42 | 2154 | 0.01 | | |
| Total | 40.46 | 2159 | | | |

**Table S21 | Estimated value of difference (EVD) across gradients and groups.**

| comparison | lower CL | estimate | upper CL | p-value |
|---|---|---|---|---|
| SC-G1 X CB-G1 | 0.0924 | 0.1172 | 0.1421 | < 0.001 |
| SC-G1 X SC-G2 | 0.1855 | 0.2103 | 0.2352 | < 0.001 |
| SC-G1 X CB-G2 | 0.1126 | 0.1374 | 0.1622 | < 0.001 |
| SC-G1 X SC-G3 | 0.1767 | 0.2016 | 0.2264 | < 0.001 |

| | | | | |
|---|---|---|---|---|
| SC-G1 X CB-G3 | 0.1578 | 0.1826 | 0.2074 | < 0.001 |
| CB-G1 X SC-G2 | 0.0683 | 0.0931 | 0.1179 | < 0.001 |
| CB-G1 X CB-G2 | -0.0047 | 0.0202 | 0.045 | 0.1886 |
| CB-G1 X SC-G3 | 0.0595 | 0.0843 | 0.1092 | < 0.001 |
| CB-G1 X CB-G3 | 0.0405 | 0.0653 | 0.0902 | < 0.001 |
| SC-G2 X CB-G2 | -0.0978 | -0.0729 | -0.0481 | < 0.001 |
| SC-G2 X SC-G3 | -0.0336 | -0.0088 | 0.0161 | 0.9164 |
| SC-G2 X CB-G3 | -0.0526 | -0.0277 | -0.0029 | 0.0181 |
| CB-G2 X SC-G3 | 0.0394 | 0.0642 | 0.089 | < 0.001 |
| CB-G2 X CB-G3 | 0.0204 | 0.0452 | 0.07 | < 0.001 |
| SC-G3 X CB-G3 | -0.0438 | -0.019 | 0.0058 | 0.2471 |

**Table S22 | Regions that showed significant differences in structure-function coupling.**

| region | hemi | yeo | SC | CB | Difference | p (uncorr) | p (corr) |
|---|---|---|---|---|---|---|---|
| V2 | L | VIS | -0.475 | 0.328 | -0.803 | 0.001 | 0.043 |
| V3 | L | VIS | -0.505 | 0.236 | -0.741 | 0.002 | 0.044 |
| IPS1 | L | DAN | -0.405 | 0.398 | -0.803 | 0.001 | 0.043 |
| FFC | L | VIS | -0.42 | 0.315 | -0.734 | 0.003 | 0.045 |
| STV | L | DMN | 0.428 | -0.333 | 0.761 | 0.002 | 0.044 |
| 8BL | L | DMN | 0.438 | -0.31 | 0.748 | 0.002 | 0.044 |
| 47s | L | DMN | 0.476 | -0.542 | 1.019 | 0 | 0.004 |
| Pir | L | LIN | 0.184 | -0.514 | 0.699 | 0.003 | 0.047 |
| STSvp | L | DMN | 0.17 | -0.675 | 0.845 | 0 | 0.014 |
| VMV1 | L | VIS | -0.488 | 0.244 | -0.732 | 0.002 | 0.044 |
| VMV3 | L | VIS | -0.469 | 0.278 | -0.747 | 0.002 | 0.044 |
| V3CD | L | VIS | -0.436 | 0.329 | -0.765 | 0.002 | 0.044 |
| VVC | L | VIS | -0.498 | 0.345 | -0.843 | 0 | 0.031 |
| STSva | L | DMN | 0.095 | -0.582 | 0.677 | 0.003 | 0.046 |
| V2 | R | VIS | -0.58 | 0.41 | -0.99 | 0 | 0.004 |
| V3 | R | VIS | -0.558 | 0.216 | -0.774 | 0.001 | 0.043 |
| V4 | R | VIS | -0.521 | 0.183 | -0.704 | 0.003 | 0.046 |
| IPS1 | R | DAN | -0.437 | 0.352 | -0.789 | 0.001 | 0.044 |
| FFC | R | VIS | -0.56 | 0.145 | -0.705 | 0.003 | 0.044 |
| V3B | R | VIS | -0.511 | 0.328 | -0.839 | 0 | 0.031 |
| PIT | R | VIS | -0.478 | 0.294 | -0.771 | 0.001 | 0.044 |
| ProS | R | VIS | -0.088 | 0.603 | -0.691 | 0.002 | 0.044 |
| V3CD | R | VIS | -0.452 | 0.312 | -0.764 | 0.002 | 0.044 |

| | | | | | | | |
|---|---|---|---|---|---|---|---|
| VVC | R | VIS | -0.522 | 0.235 | -0.757 | 0.001 | 0.044 |
| STSva | R | DMN | 0.302 | -0.426 | 0.728 | 0.003 | 0.046 |
